# Markov models bridge behavioral strategies and circuit principles facilitating thermoregulation

**DOI:** 10.1101/2025.03.17.643749

**Authors:** Kaarthik Abhinav Balakrishnan, Martin Haesemeyer

## Abstract

Behavioral thermoregulation is critical for survival across animals, including endothermic mammals. However, we do not understand how neural circuits control navigation towards preferred temperatures. Zebrafish exclusively regulate body temperature via behavior, making them ideal for studying thermal navigation. Here, we combine behavioral analysis, machine learning and calcium imaging to understand how larval zebrafish seek out preferred temperatures within thermal gradients. By developing a stimulus-controlled Markov model of thermal navigation we find that hot avoidance largely relies on the modulation of individual swim decisions. The avoidance of cold temperatures, a particular challenge in ectotherms, however relies on a deliberate strategy combining gradient alignment and directed reversals. Calcium imaging identified neurons within the medulla encoding thermal stimuli that form a place-code like representation of the gradient. Our findings establish a key link between neural activity and thermoregulatory behavior, elucidating the neural basis of how animals seek out preferred temperatures.

## Introduction

Biological processes are optimized for specific temperatures. Deviations from these temperatures can have severe or fatal consequences ^1^–4. Organisms therefore evolved mechanisms to minimize such deviations ^5^–7 including the ability to seek out preferred temperatures through thermoregulatory behaviors like thermal gradient navigation or sunbathing^8^. Endotherms including mammals and birds additionally regulate body temperature using autonomous mechanisms, such as sweating, shivering, and modulating blood flow ^9,10^. Despite this ability, thermoregulatory behaviors are conserved in endotherms because these are more energy efficient than autonomous regulation ^11,12^. Therefore, mammals will readily navigate thermal gradients to seek out preferred temperatures ^13–19^. Ectotherms lack internal physiological mechanisms for temperature regulation and exclusively rely on behavior, such as exploring their environment to find optimal temperatures ^9,20^. Despite the universal importance and high conservation, thermoregulatory behavior and its neural basis remain poorly understood in vertebrates ^21,22^.

Here we combined behavioral recording, modeling, and functional calcium imaging in larval zebrafish to gain insight into how vertebrates seek out preferred temperatures through the interaction of two competing drives: the avoidance of hot and cold temperatures, respectively. As ectotherms, larval zebrafish seek out preferred temperatures to thermoregulate and due to their small size, they are largely at equilibrium with the temperature in their environment. Previous work in larval zebrafish uncovered behavioral features and neural circuits of heat avoidance ^23–25^, how swim kinematics and turn sequencing change with temperature ^26,27^ and demonstrated that structures implicated in mammalian thermoregulation have conserved roles in zebrafish^28^. However, none of these studies addressed how larval zebrafish seek out a preferred temperature. This goal directed behavior crucially requires a change in valence attached to thermal stimuli. While heating stimuli signal worsening conditions above the preference (“worsening context”^28^), they signal improving conditions below. This change in the valence of stimuli is a critical feature of homeostatic regulation. We therefore focused on comparing behavioral mechanisms of hot and cold avoidance and how neurons within the medulla of larval zebrafish encode features of temperature gradients to enable this switch in stimulus valence.

Specifically, we characterized zebrafish behavior across a range of temperatures, focusing on their navigation strategies under different conditions, including uniform (“constant”) temperatures and temperature gradients. Comparison of movement strategies under uniform and gradient conditions identified specific behavioral modes that are critical for cold avoidance but expendable for hot avoidance. To capture this structure, we fit a “Navigation model”, combining a Markov model with stimulus-dependent transitions and stimulus-dependent behavioral emission models. Simulations revealed that this model captures both hot and cold-avoidance behavior. Using functional calcium imaging we identified the representation of hot and cold temperatures within the zebrafish medulla. These form a “place code” within temperature gradients that underlies the swim-mode transitions in the Navigation model.

## Results

### A comparative approach to identify thermoregulatory strategies

To investigate how larval zebrafish navigate temperature gradients to thermoregulate, we designed a gradient chamber similar to Le Goc ^26^, see Methods (Figure 1A). Individual larvae swam in two 200 mm long and 50 mm wide lanes within a machined aluminum chamber. We used a custom-programmed temperature controller to establish linear temperature gradients or uniform temperature profiles (“constant temperature”) within the chamber by using Peltier elements for heating and cooling. The position and heading of zebrafish were tracked under white-light illumination at 100 Hz while they explored the chamber (Figure 1B). This allowed us to extract swim kinematics (Figure 1B) such as changes in heading angles (turn angles), the waiting time between consecutive swim bouts (interbout-interval) as well as the total displacement for each swim (bout displacement).

**Figure 1.**
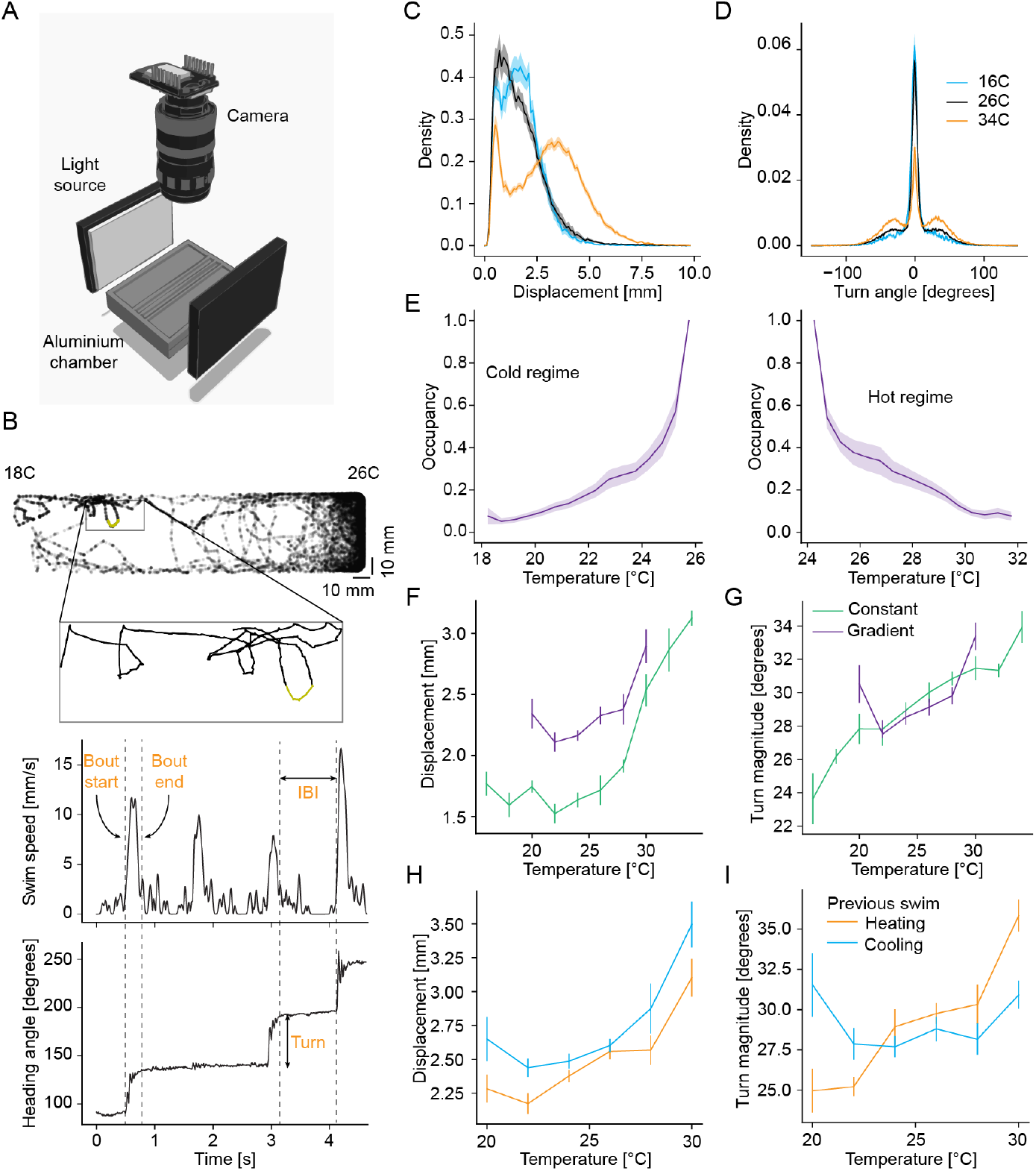
Comparison of bout parameters of larval zebrafish across experimental conditions. **A** Illustration of the behavioral setup to track fish in a temperature controlled aluminium chamber. **B** Example tracking of larval zebrafish in an experiment with a highlighted section of the trajectory shown in a zoomed inset; the fish movement in that trajectory (top) is characterized using instantaneous swim speed (middle) and heading angle (bottom), with the definition of bout starts, bout ends, inter-bout intervals and turn angle indicated on the plot. **C** Example distribution of swim displacements (in mm) at three different temperatures averaged over all corresponding constant temperature experiments (blue-16 °C, black-26 °C, orange-34 °C). **D** Example distribution of turn angles (in degrees) for the same temperatures as (C). **E** Occupancy of larval zebrafish in the chamber averaged across all experiments with a temperature gradient in the Cold regime and Hot regime. **F** Comparison of median swim displacements (in mm) of larval zebrafish with respect to temperature in constant temperature (green) and gradient (purple) experiments. **G** Comparison of median turn magnitudes (in degrees) of larval zebrafish with respect to temperature in constant temperature (green) and gradient (purple) experiments. **H** Comparison of median swim displacements of larval zebrafish at each temperature for different contexts in gradient experiments-orange-heating context, blue-cooling context. **I** Comparison of median absolute turn magnitude of larval zebrafish at each temperature for different contexts in gradient experiments-orange-heating context, blue-cooling context. All error bars and shaded error regions are bootstrap standard errors across experiments.

As a baseline for our subsequent analysis of behavior within the gradient, we isolated the effect of absolute temperature on swim kinematics within our setup. As observed by Le Goc ^26^, the frequency and vigor of swims increased overall with increasing temperature. Specifically, the interbout interval (IBI) decreased with increasing temperature (Figure S1A), while the displacements and turn magnitudes of swims increased with increasing temperatures (Figure 1C-D). The fraction of straight swims (turn magnitude smaller than 5 ^*°*^, see Methods) especially decreased as temperatures increased (Figure 1D, reduction in central peak). These effects could be a mixture of changes in neural control as well as direct temperature effects on muscle physiology of ectothermic fish. However, it is interesting to note that swim displacements did increase again at the coldest temperatures (Figure 1C shift in distribution maximum at 16 °C) indicating that there was no general impairment of movement. Overall, these results suggest that larval zebrafish need an efficient strategy to escape from the cold as their movement repertoire was more limited under these conditions.

Despite these challenges, larval zebrafish readily avoided cold temperatures when placed into a gradient ranging from 18 °C to 26 °C (slope of 0.04 °C/mm) (Figure 1E). In fact, zebrafish avoided cold temperatures at least as well as they avoided hot temperatures within a gradient ranging from 24 °C to 32 °C on the hot side (matched slope of gradient, Figure 1F) despite their reduced movement at colder temperatures. In constant temperature experiments, fish would generally accumulate near the edges of the chamber (thigmotaxis, Figure S1B); and in the absence of a gradient (uniform constant temperature), position along the chamber axis had very little influence on swim kinematics (Figure S1C-E). To better understand the effect of the gradient on swim kinematics we compared behavior kinematics at matched temperatures within the gradient to those of the constant temperature experiments. The main difference between gradient and constant experiments was that, within the gradient, changes in position during swim bouts were coupled to changes in temperature (Figure S1F-G). This change in stimulus context increased overall swim vigor. Displacements were higher within the gradient (Figure 1H), while interbout intervals were shorter, especially at temperatures below 24 °C C (Figure S1H). The enhancement of displacements at lower temperatures within the gradient compared to constant temperatures indicated that at least part of the reduction in swim vigor at colder temperatures could be compensated if swims served a behavioral goal (i.e., reaching the preferred temperature^29^). Turn angles on the other hand were largely comparable between the two experiment types (Figure 1I). These changes were also reflected in an influence of the change in temperature experienced during previous swim bouts on subsequent swim kinematics (Figure S1I-K).

To analyze these effects in the context of navigation, we split swim bouts according to whether the previous swim led to a temperature increase (“Heating”) or decrease (“Cooling”) using a threshold of 0.052 °C/bout as the cutoff which corresponded to the upper and lower quartiles of temperature changes experienced by the larvae during the experiments (Figure S1F). Across the entire temperature range tested, swim displacements were larger after temperature decreases than after temperature increases (Figure 1H). Interbout intervals on the other hand were not influenced by the direction of temperature change, except below 24 °C where cooling stimuli shortened the interbout interval (Figure S1L). Similarly to the effect of constant temperature, modulation of swim vigor by changes in temperature appeared counterproductive for cold avoidance: larval zebrafish swam further and more often on trajectories away from the preference, towards colder waters. On the other hand, the magnitude of turning was modulated by context rather than the absolute direction of temperature change (Figure 1I). Specifically, above 26 °C and below 23 °C, zebrafish increased the magnitude of turns during worsening contexts: heating stimuli increased turning in warm temperatures as observed in Palieri ^28^, while temperature decreases led to an increase in turning in cold temperatures.

In summary, larval zebrafish readily avoided both hot and cold temperatures. However, absolute temperature appeared to change behavior in a manner that opposed cold avoidance. The same was true for the influence of Heating/Cooling stimuli on swim vigor which favored accumulation at colder temperatures. Given the prominent cold avoidance behavior, these effects were likely offset by a more deliberate strategy of avoiding cold. This can be partially attributed to the worsening temperature context modulating turn magnitude to a similar degree at cold temperatures as it does at hot temperatures. This modulation could reorient zebrafish when swimming away from the preferred temperature range.

### Modulation of individual swim kinematics explains hot avoidance

We performed simulations to understand the relationship of the modulation in bout kinematics to gradient navigation in both the hot (24-32 °C gradient) and cold (18-26 °C gradient) regimes. Initially, we created a non-parametric model of fish behavior by binning the swim data obtained during the constant temperature and gradient experiments according to absolute temperature, as well as changes in temperature experienced during the previous swim bout. Such a model can accurately capture the true swim repertoire of larval zebrafish. However, it lacks a functional form to encapsulate stimulus processing. As observed by Le Goc ^26^ the exclusive modulation of swim kinematics by absolute temperature led to an accumulation of simulated fish at the cold edges of the gradient, both in the hot and cold regime (Figure 2A-C). This partly explained hot avoidance but failed to explain how larval zebrafish avoid cold temperatures. To quantify how well a given model captures real fish behavior, we computed the Kullback-Leibler (KL) divergence between the observed gradient distribution and distributions resulting from simulations. As expected, based on the slow accumulation on the cold side, the constant temperature simulation improved hot avoidance compared to random behavior (uniform distribution) but was worse than random behavior for avoiding cold (Figure 2B-C).

**Figure 2.**
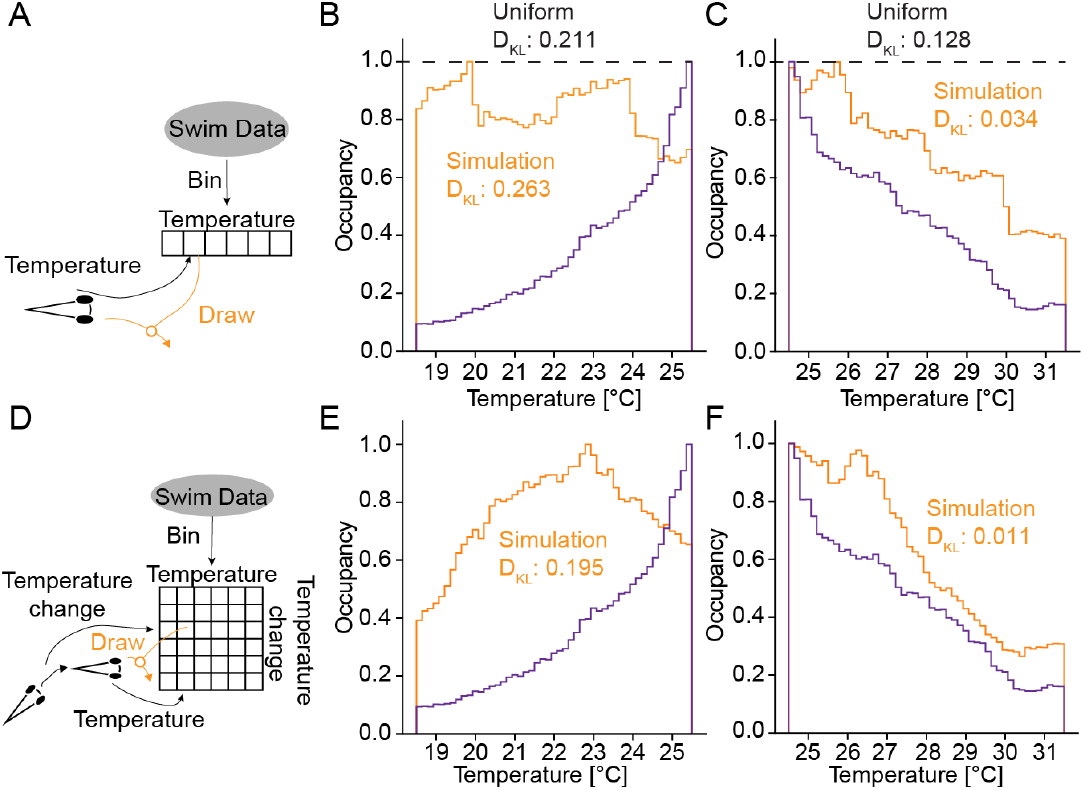
Simulation of non-parametric model of navigation based on experimental distribution of bout parameters shows cold avoidance. **A** Schematic of simulation of fish in a virtual gradient with bout movements picked from binned values of constant temperature experiments. **B-C** Comparison of occupancy of fish in gradient temperature experiments (purple) and simulations using the non-parametric model (orange) binned from constant temperature experiments in the cold (B) and hot (C) regime. Black dashed line indicates uniform distribution of a randomly moving particle. *DKL* indicates KL divergence between simulations as well as the uniform distribution and the fish data. **D** Schematic of simulation of fish in a virtual gradient with bout movements picked from gradient temperature experiments, binned according to temperature and temperature change. **E-F** Same as (B-C) using the nonparametric temperature/temperature change simulation.

Absolute temperature could inform zebrafish about how far away they are from the preference but not whether they are swimming towards it (improving context) or away from it (worsening context). We therefore expanded our model and included the modulation of swim kinematics by changes in temperature (Figure 2D). This model led to a dramatic improvement in hot avoidance: The resulting distribution mimicked the true fish distribution closely as evidenced by a small KL divergence (Figure 2F). Including temperature changes in the model also improved cold avoidance, bringing the distribution of the simulated fish closer to the real fish distribution compared to random behavior (Figure 2E). This indicated that hot avoidance could be explained by considering swim-by-swim modulations of kinematics, which were driven by absolute temperature and changes in temperature. Cold avoidance on the other hand likely relied on additional behavioral modulation.

Based on the outcomes of the simulations, we postulate that zebrafish navigate their environment using different strategies for cold temperatures versus hot temperatures. They likely followed a more deliberate and efficient navigation strategy in the cold, since they tended to use fewer and slower swims to escape these temperatures. In contrast, in warmer temperatures, they exhibited more frequent swims of larger displacement, which indicates a strategy akin to a biased random walk, where sensory stimuli modulated behavior on a swim-by-swim basis.

### Larval zebrafish combine gradient alignment and directed reversals to avoid cold

Within the gradient we observed strong autocorrelations between the displacements of successive swims indicating that larval zebrafish maintain swim displacements over longer timescales. These autocorrelations were larger at higher temperatures (Figure 3A). As observed in Palieri ^28^, temperature increases led to a larger correlation in successive turn angles, likely aiding reorientation during hot avoidance (Figure 3B). However, this modulation was not observed at colder temperatures (Figure 3B). In constant temperature experiments (as well as for swims within the gradient without a preceding change in temperature), auto-correlations were modulated by temperature. Correlations of successive displacements increased with increasing temperature while correlations of successive turns decreased (Figure S2A-B). Because of the modulation of correlations by the stimulus we wondered whether larval zebrafish used longer-term behavioral strategies to avoid cold temperatures. Such strategies would likely be necessary to overcome problems with reduced movement frequency and vigor in the cold.

**Figure 3.**
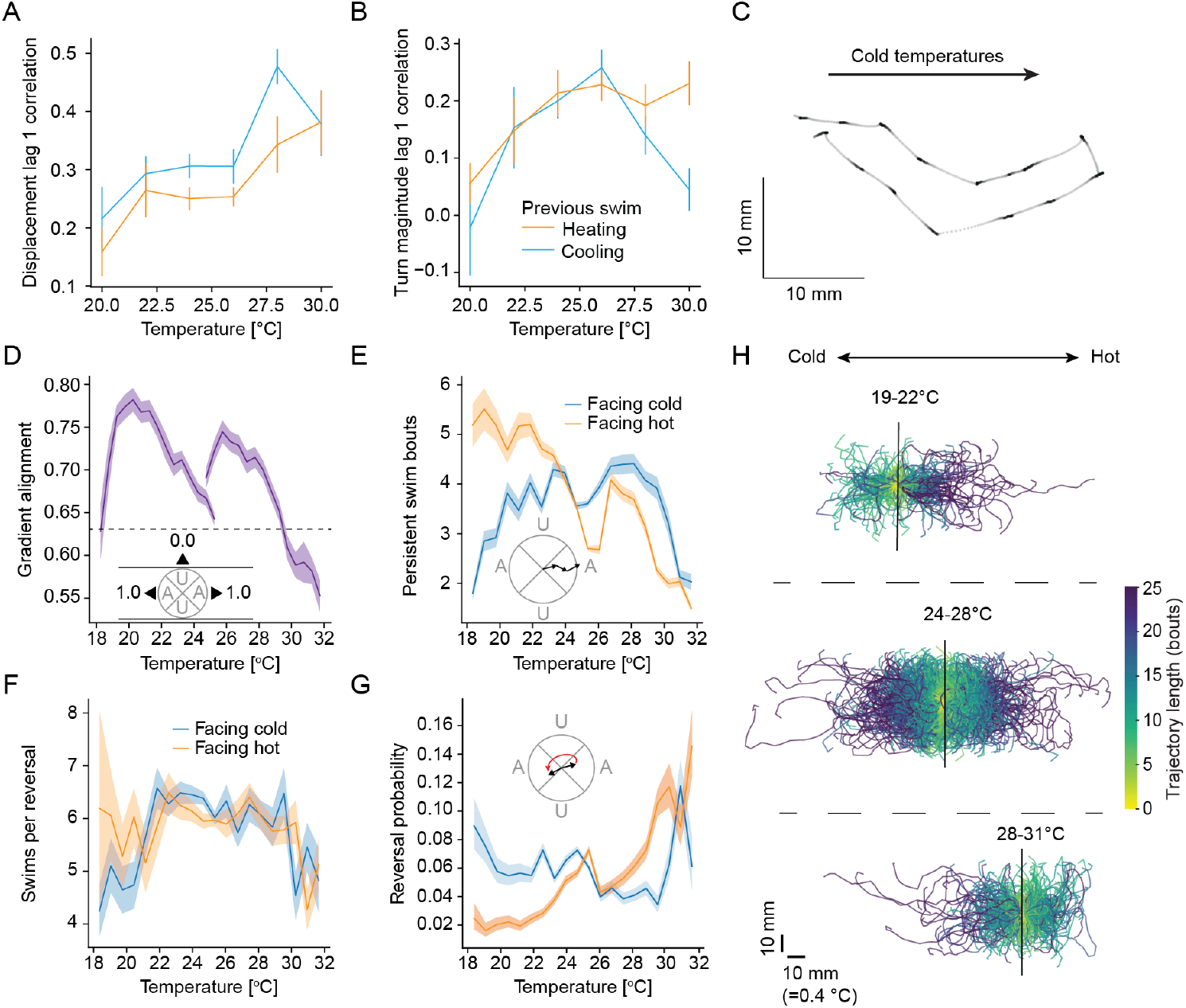
Larval zebrafish modulate gradient alignment, persistent swims and reversals to aid navigation. **A** Autocorrelation of displacement in consecutive swims at different temperatures in different contexts-heating (orange) and cooling (blue). **B** Autocorrelation of absolute turn magnitude in consecutive swims at different temperatures in different contexts-heating (orange) and cooling (blue). **C** Example experimental trajectory of cold avoidance in larval zebrafish showing gradient aligned swimming and a reversal maneuver. **D** Alignment of larval zebrafish with the gradient direction at different temperatures in hot and cold regimes compared to a random gradient alignment (black dotted line); inset figure shows the definition of the cones of alignment with gradient direction. **E** Average number of swim bouts in persistent trajectories starting at different temperatures under different contexts-facing hot direction (orange) and facing cold direction (blue); inset figure illustrates the definition of a persistent trajectory. **F** Average number of swim bouts in reversal manuevers starting at different temperatures under different contexts-facing hot direction (orange) and facing cold direction (blue). **G** Probability of initiating a reversal on trajectories starting at different temperatures under different contexts-facing hot direction (orange) and facing cold direction (blue); inset figure shows the definition of a reversal trajectory. **H** Examples of trajectories until initiation of a reversal from gradient experiments with the larval zebrafish starting at different temperatures (black lines); colorbar represents number of swim bouts until reversal start. All error bars and shaded error regions are bootstrap standard errors across experiments.

Close observation of trajectories during cold avoidance suggested that larval zebrafish often swam along the gradient direction in the cold. Furthermore, when they swam away from their preference (to even colder temperatures, worsening context) zebrafish appeared to terminate these aligned movements with sharp reversals (Figure 3C). These reversals are reminiscent of C. elegans pirouettes during hot avoidance ^30^. To analyze these movement modes, we defined a gradient alignment metric. For each swim, we computed the absolute cosine of the angle of the swim vector (from start of swim to end) relative to the gradient direction. This “alignment metric” is 1 if the swim is perfectly aligned to the gradient, and 0 if the swim is perpendicular.

Within the gradient, the alignment metric showed a clear temperature dependence: at colder temperatures, swims were on average more aligned to the gradient than chance, while they were less aligned than chance in hot temperatures (Figure 3D). In constant temperature experiments alignment was independent of position (Figure S2C) but modulated by temperature, however to a lesser extent than in the gradient (Figure S2D). Gradient alignment was therefore likely driven by a combination of thermal and nonthermal features such as visual landmarks. The stronger modulation in the gradient indicates that larval zebrafish detected the thermal gradient direction and actively aligned to it in the cold; in hot conditions they conversely used gradient information to suppress gradient alignment. To capture this phenomenon, we divided swims into “aligned swims” (absolute cosine *>* 0.71) and “non-aligned swims” (absolute cosine *<* 0.71). This allowed us to investigate if changes in alignment are dependent on worsening vs. improving context in addition to absolute temperature. To this end, we defined persistence length as the number of successive aligned swims. The average persistence length was highest (*>* 5 bouts) in the cold when context was improving (facing hot, Figure 3E)). Persistence length was strongly reduced during worsening context in the cold (Figure 3E). As expected from the overall effect of temperature on alignment, modulation of persistence length was considerably smaller above the preferred temperature. This indicated an efficient strategy centered on maintaining or breaking gradient alignment in the cold based on context. As expected, persistence length was not modulated by direction in constant temperature, but was overall shorter in higher temperatures, likely due to the overall increase in turn magnitude as temperatures increase (Figure S2E-F).

To understand whether larval zebrafish reverse direction during worsening contexts in the cold, we subsequently defined reversals as swim sequences that take larval zebrafish from one aligned direction (e.g., facing towards the cold) to the opposite (facing towards warmth). Intriguingly, the average length of reversals was around 6 swim bouts irrespective of temperature (Figure 3F) despite the overall increase in turn magnitude at higher temperatures. Furthermore, across reversals 85 % of bouts turned in the same direction, regardless of temperature or context (Figure S2G). This indicates that larval zebrafish executed reversal maneuvers in a ballistic fashion, possibly under dedicated neural control. Analyzing reversals by worsening or improving context revealed the opposite dependency compared to persistence. We found that the probability to initiate a reversal maneuver was essentially constant whenever larvae were facing colder temperatures. However, reversals were strongly suppressed at cold temperatures when zebrafish were facing towards their preferred temperature (Figure 3G). At hot temperatures, the probability of reversals was similarly increased during worsening context, however, the relative increase was smaller than in the cold. The smaller contrast in reversal initiation at hot temperatures was expected considering the fact that a simple swim-by-swim simulation was nearly sufficient in explaining hot avoidance. The difference in reversal probabilities led to a clear modulation in trajectory length until reversals occurred based on position and direction (Figure 3H). In the cold, reversals were often initiated within less than ten swim bouts when zebrafish were swimming away from the preference while fish often maintained direction for more then twenty bouts when swimming towards warm temperatures (Figure 3H, top). Near the preferred temperature, direction within the gradient did not influence the length of trajectories until reversals occurred (Figure 3H, middle). At hot temperatures the amount of swim bouts until reversal was again reduced for trajectories away from the preference (Figure 3H, bottom). In constant temperature experiments, reversal probability was slightly modulated by chamber position and was maximal for intermediate temperatures (Figure S2H-I). However, modulation by swim direction within the chamber was small and opposite to what was observed within the gradient, strongly arguing that it was the change in temperature that modulated reversal probabilities. In line with this, starting from the same chamber positions as in the gradient experiments, trajectories until reversal were not modulated by direction in constant temperature experiments (comparing Figure S2J with 3H).

Taken together, these results suggest an asymmetry in navigation at cold versus hot temperatures. Zebrafish biased their swim trajectories in colder temperatures such that they initiated reversals more easily during worsening context, and initiated more persistent, gradient aligned, trajectories in improving contexts. On the other hand, in hot temperatures these longer-timescale swim modes were modulated less by worsening versus improving context.

### A GLM driven Markov model captures gradient behavior

To understand how larval zebrafish thermoregulate by avoiding cold and hot temperatures, we sought to capture the relevant behavioral features in a quantitative, parametric “Navigation model” (Figure 4A). We designed a Markov model to capture swim modes with transitions dictated by the stimulus (temperature *T* and temperature change across the previous swim Δ*T*) through a generalized linear model (GLM). Based on our observations we defined three swim modes: A “Reversal mode” capturing reversal maneuvers, a “Persistent mode” capturing swims with a persistent direction and a “General mode” for all other swims (see Methods). While the consistency of length and handedness of reversals suggested that these may be fixed action sequences, we nonetheless defined the outputs within each mode to be individual swims rather than full sequences. Firstly, this appeared prudent since we do not have direct evidence that zebrafish initiate compound reversals; secondly, it allowed us to define the model in terms of the smallest movement unit generally used in the field, individual swim bouts. For each swim mode we fit separate output models using GLMs that generate interbout intervals (modeled as Gamma distributions), displacements (modeled as Gamma distributions) and turns (modeled as Gaussian-Gamma-mixture models). Similar to the GLM capturing transitions between swim-modes, the output GLMs used the stimulus to compute behavior. In addition, output GLMs also included previous behavior as inputs to capture observed auto-correlations (see Methods and Supplemental Table 1). Crucially, the preference presents a discontinuity for the control of behavior, i.e., the worsening context is defined by increasing temperatures above but decreasing temperatures below the preference. Therefore, linear processing of the stimulus cannot account for both hot and cold avoidance and linear processing would not be able to capture the observed modulation of turn magnitude according to context (Figure 1I). We therefore opted for 2nd order GLMs including interaction terms in our output models. In the following, we use the term “Navigation model” to refer to the entire model consisting of the Markov model governing transitions between swim modes and the mode-specific emission models that generate swim kinematics.

**Figure 4.**
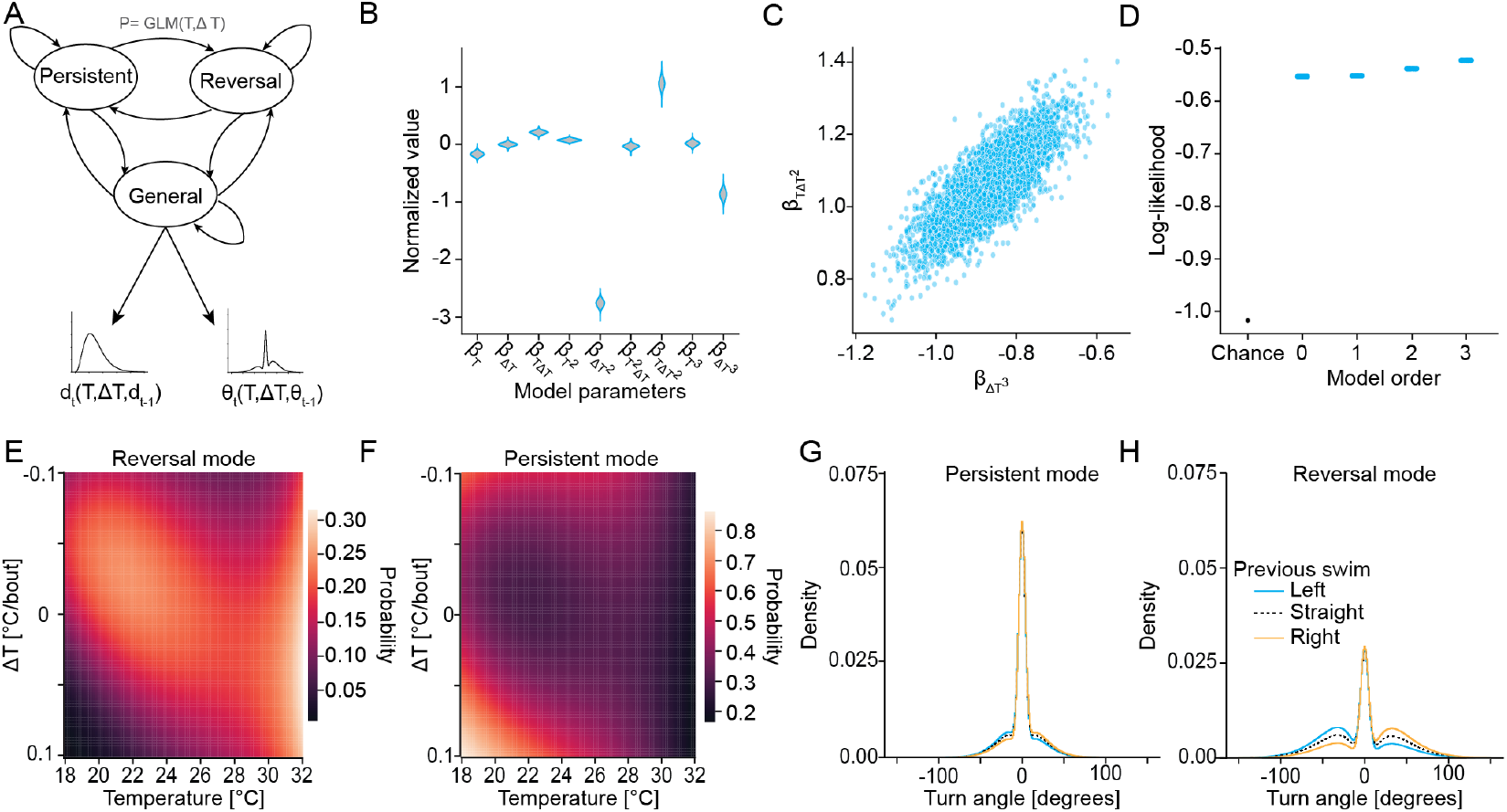
Modeling larval zebrafish behavior using Markov models highlights the effect of temperature and change in temperature (context) on swim modes. **A** Schematic of Markov model to describe larval zebrafish movements, with transition probabilities between different modes being controlled by a GLM that depends on previous temperature and change in temperature; followed by emission probabilities controlled by a GLM that depends on temperature, previous temperature and behavioral history (previous displacement for displacement emissions and previous turn for turn emissions). **B** Example distribution of GLM parameters, normalized according to the standard deviation of the quantity the parameter acts on, obtained from Monte Carlo fits for the reversal-to-reversal transition probability. Note that each violin plot encompasses 4000 draws from the posterior distribution. **C** Scatter plot of GLM parameters related to *T* Δ*T* ^2^ and Δ*T* ^3^ showing a strong correlation (*r* = 0.8). **D** Comparison of goodness of fit of transition models of different orders using log-likelihood of model predictions on test-data, with the chance model which uses the overall data distribution for predictions. **E** Heatmap of steady state probabilities of the reversal mode depending on temperature and change in temperature for the 3rd order model. **F** Heatmap of steady state probabilities of the persistent mode depending on temperature and change in temperature for the 3rd order model. **G** Dependence of emission of turn angle on the previous turn in the persistent mode - previous left turn (blue), previous straight swim (black dotted) and previous right turn (orange). **H** Same as (G) for reversal mode.

We used Markov-Chain-Monte Carlo (specifically, Hamiltonian Monte Carlo ^31^) to fit all transition and output parameters of the Navigation model. The advantage of this approach was that all parameters were sampled from a joint posterior distribution ^31^,32. This allowed us to perform all model tests and simulations using draws from this distribution rather than relying on a maximum likelihood estimate. Furthermore, pairwise correlations between parameters can reveal flexible directions in model space, i.e. fixed relationships between parameters while individual parameter values can fluctuate without impacting model performance. As an example, the transition GLM from the Reversal mode back to the Reversal mode (maintenance of reversal) heavily relied on higher-order interactions of current temperature and the change in temperature during the previous bout (Figure 4B). This was expected, since reversals depend on worsening vs. improving context, which in turn depends on an interaction of where the fish is within the gradient and the direction in which it is swimming. In this GLM, parameter values for the interaction between temperature and the squared change in temperature *T* Δ*T* ^2^ strongly correlated (*r* = 0.8) with parameter values of Δ*T* ^3^ (Figure 4C). This correlation was not the result of correlations in the corresponding inputs which were only nonlinearly related (*r* = *−*0.05) (Figure S3A).

This indicates that the ratio of these parameters was largely fixed, while individual values are flexible. To assess the good-ness of fit for our model, we split the experimental data in a training set (80% of all swims) and a test set (20% of all swims). We subsequently computed the log-likelihood of model predictions of swim modes (transition model) as well as kinematic features (output models) on the test data and compared it to a “chance model” that uses the overall data distribution for predictions (see Methods). In all cases, the log-likelihood of the fit model surpassed that of the chance model (Figure S3B). For the transition model, we considered increasing orders of the GLM; a 0th order model predicts constant transition probabilities (as given across the entire dataset), a 1st order model allows for linear dependencies on the stimulus (temperature and temperature change), while 2nd and 3rd order models include interaction terms. The largest jump in log-likelihood of the test data under the model occurred from a chance model (assume that each mode occurs at its given probability irrespective of the preceding mode) to the 0th order model (assume fixed transitions). This simply indicated that swim modes indeed followed a Markov process. From 0th to 3rd order we observed a steady, significant, increase in log-likelihood (Figure 4D). GLM orders higher than 3rd were not tested due to the explosion in size of the parameter space. We therefore chose the 3rd order model for further analysis.

We computed the steady-state probabilities predicted for each swim mode from the transition matrix of the Markov model. We specifically analyzed the steadystate probabilities in relation to the temperature change in the previous bout as well as the current temperature. This revealed how larval zebrafish distributed swim modes within the gradient (Figure 4E-F). At temperatures below 22 °C the Reversal mode probability was strongly influenced by context (Figure 4E). While the Reversal mode was almost absent during improving context it was prevalent during worsening context (decreasing temperatures). Close to the preference (25 °C -28 °C), the asymmetry in the influence of context was considerably milder while it increased again as zebrafish entered hot temperatures (Figure 4E). Above 29 °C the

Reversal mode was favored in the worsening context compared to improving context. However, as observed on the raw behavior data the contrast between improving and worsening context was stronger for cold than hot temperatures. This was even clearer for the Persistent mode (persistent alignment to the gradient direction) which was strongly favored during improving context in cold temperatures but which was largely absent at hot temperatures irrespective of the context (Figure 4F). Instead, we observed that zebrafish were more likely to be in the General mode at hot temperatures (Figure S3C). This further underlined that movement relative to the gradient direction played a smaller role during hot avoidance while zebrafish instead relied on biased individual movements to escape heat (swim-by-swim modulation). Since probabilities across all modes sum to one, steady-state probabilities of the General mode largely reflected overall decreases in reversal and persistent probabilities. As such, the General mode was most prevalent at temperatures above 28 °C (Figure S3C).

The emission models reflected the expected behavior within each mode. The Persistent mode was dominated by straight swims with smaller angle turns (Figure 4G). There was a slight dependence of the current turn direction on the direction of the previous turn, a general feature of larval zebrafish movement^33^. Reversals on the other hand were dominated by large angle turns. Furthermore, the previous turn direction strongly influenced the direction of the current turn (Figure 4H) as evidenced by the asymmetry of turn direction distributions induced by the model. This was expected, since reversals are characterized by larval zebrafish stringing together turns of the same direction (Figure S2G). Displacements on the other hand were largely independent of the swim mode but overall strongly influenced by the displacement of the previous swim bout (Figure S3D-E).

In summary, the Navigation model allowed us to capture and characterize the dependence of swim modes as well as individual behaviors on the temperature stimulus and past movement history. This revealed important dependencies of swim mode probabilities on the temperature stimulus which likely underlie the ability of larval zebrafish to efficiently navigate away from the cold while swim modes are less important for hot avoidance.

### Cold avoidance requires a non-linear transition model

After establishing a navigation model that incorporated observed behavioral modes and mode-specific generation of behavior, we set out to test whether this model could be used to simulate navigation of temperature gradients. We specifically focused on assessing the importance of the structure of the Markov model since understanding how the stimulus governs swim mode transitions makes predictions about neural processes that control swim modes. To this end, we used the navigation model to simulate exploration within virtual temperature gradients matching the experimental gradients (Figure 5A, see Methods). In all simulations the emission models were the same, having a quadratic (2nd order) dependence on temperature and temperature-change together with relevant behavioral history and interaction terms. Instead, we varied the order of the transition model to assess how this would affect gradient navigation performance.

**Figure 5:**
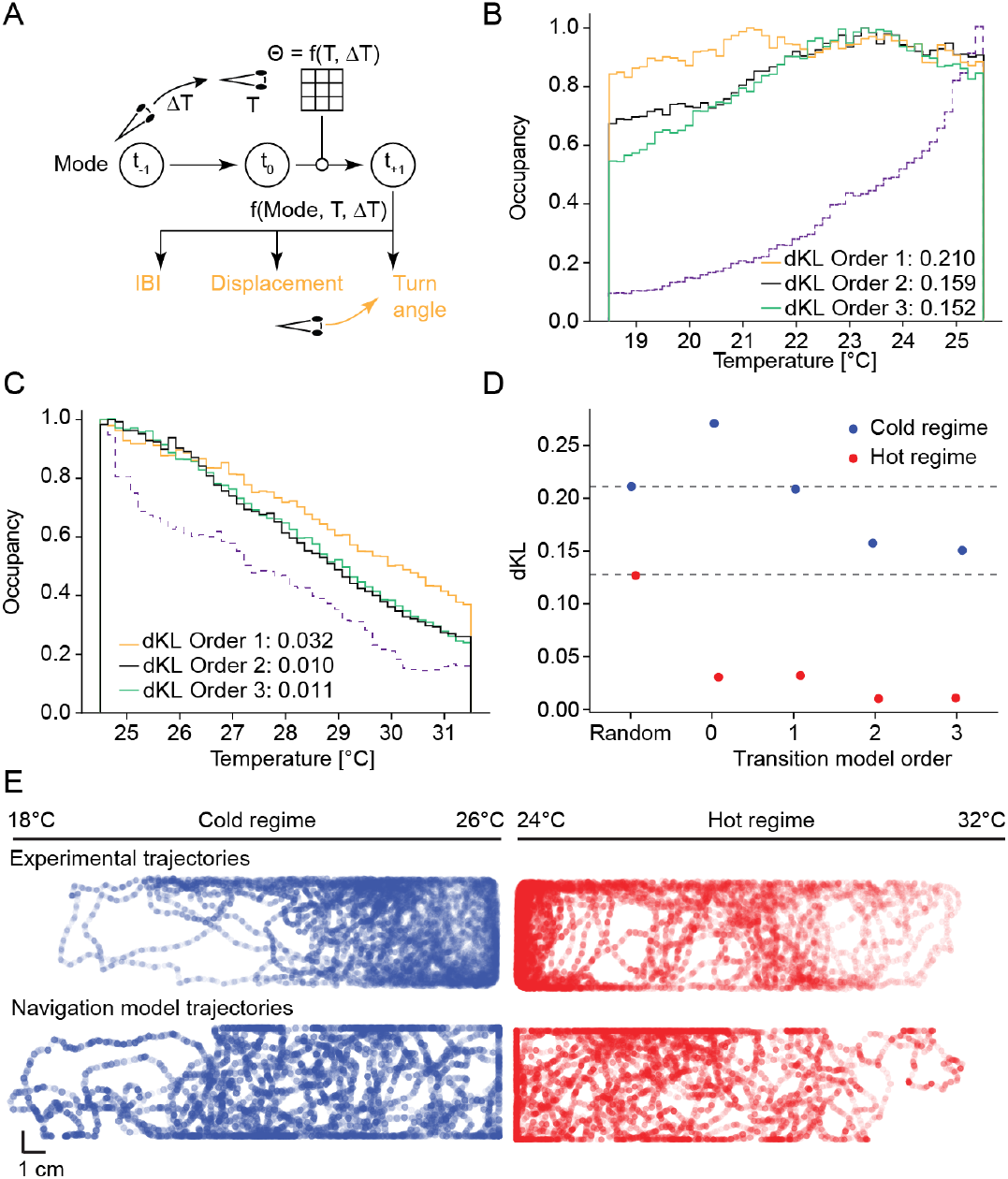
Simulation of fish behavior using a parametric Navigation model reveals the importance of nonlinear control of transitions for cold avoidance. **A** Schematic of simulation of larval zebrafish in a virtual gradient using the Navigation model to generate a new state, followed by emission of bout parameters-IBI, displacement and turn angle. **B** Comparison of occupancy of fish (purple dotted) in the cold regime and simulation using the parametric Navigation model for different orders including KL-divergences. **C** Comparison of occupancy of fish (purple dotted) in the hot regime and simulation using the parametric Navigation model for different orders including KL-divergences **D** Comparison of KL divergence for simulations using transition models of increasing order (black dotted lines represent KL divergence of a random model that does not have any temperature dependence of bout parameters) in the cold regiome (blue) and hot regime (red). **E** Comparison of tracked trajectories of larval zebrafish in a cold gradient (18 °C to 26 °C) and a hot gradient (24 °C to 32 °C) with a simulated experiment using the Navigation model with a 3rd order transition model.

A model with constant transition probabilities (0th order) led to hot avoidance (Figure S4A). For cold avoidance, however, this model was worse than a random model as it led to accumulation in the cold (Figure S4B). This mimicked the results of the nonparametric model using behavior binned according to temperature. A linear transition model (1st order) performed similarly to the constant model for hot avoidance and led to a near uniform distribution in the cold (Figure 5B-C). This suggests that a large part of hot avoidance was independent of swim modes as it was solely governed by the temperature dependence of the emission models. However, only nonlinear transition models (2nd and 3rd order) led to appreciable cold avoidance (Figure 5B), outperforming the nonparametric model of swims binned according to temperature and temperature change (Figure 2D). This strongly suggests that nonlinear processing of temperature stimuli was critical for cold avoidance and that swim modes played a large part in this behavior. Hot avoidance also improved for the nonlinear transition models, which indicates that swim modes generally contributed to navigation efficiency (Figure 5C-D) and trajectories generated by the 3rd order model mimicked behavioral trajectories of larval zebrafish (Figure 5E). The modulation of swim kinematics by gradient position and movement direction in the simulations closely matched the corresponding observations of fish navigation the thermal gradient (Figure S4C-J vs. Figures 1 and S1) with the exception of auto-correlations of successive turns (Figure S4J). This indicates that the structure of the navigation model closely captured real fish behavior. Unlike hot avoidance, the cold avoidance performance of the model did not approach that of zebrafish navigating a gradient. This was likely due to behavioral aspects not captured by the model. Firstly, the model did not have any inputs that would lead to alignment to the gradient direction; secondly, we did not treat reversals as compound behaviors which reduced the efficiency of direction reversals. These choices were made so as not to enforce any model outputs for which we currently do not fully understand the sensory or motor basis.

In summary, simulations based on the navigation model could replicate hot avoidance and large aspects of cold avoidance. By varying the order of the transition model, we could show that mode transitions are critical and have to nonlinearly depend on the stimulus. This was especially true for cold avoidance where the nonlinearity of the emission models was not enough to drive navigation but instead required nonlinearity of the transitions as well. This strongly suggested that neural circuits have to nonlinearly process the temperature stimulus to control swim modes that underlie efficient gradient navigation.

### Calcium imaging reveals stimulus segregation across seven response types within the medulla

After characterizing the behavioral algorithm of thermoregulation, we set out to identify potential neural substrates. We developed a system to present temperatures across the entire thermal range used in the behavior experiments to head-fixed larval zebrafish during functional calcium imaging (Figure 6A). Larval zebrafish expressing the calcium indicator GCaMP across all nuclei in the brain (Elavl3-H2B:GCaMP6s^34^) were embedded in low-melting point agarose within a flow chamber. A constant flow of water was provided, and an inline heater/cooler was used to adjust the temperature of the water flowing over the head while a thermistor was used to monitor the temperature at the preparation. We provided stimuli within both the absolute temperature range used in the experiments (18-32 °C), as well as temperature changes approaching those experienced by zebrafish within the gradients (Figure S5A). In previous work, we performed whole-brain imaging while presenting heating stimuli to larval zebrafish. This work suggested that neurons within the zebrafish medulla are critical to explain swim generation during heat avoidance ^25^. We therefore focused our imaging onto the medulla to understand temperature encoding across the thermal range and how thermal processing would enable the behavioral algorithm of thermoregulation.

**Figure 6.**
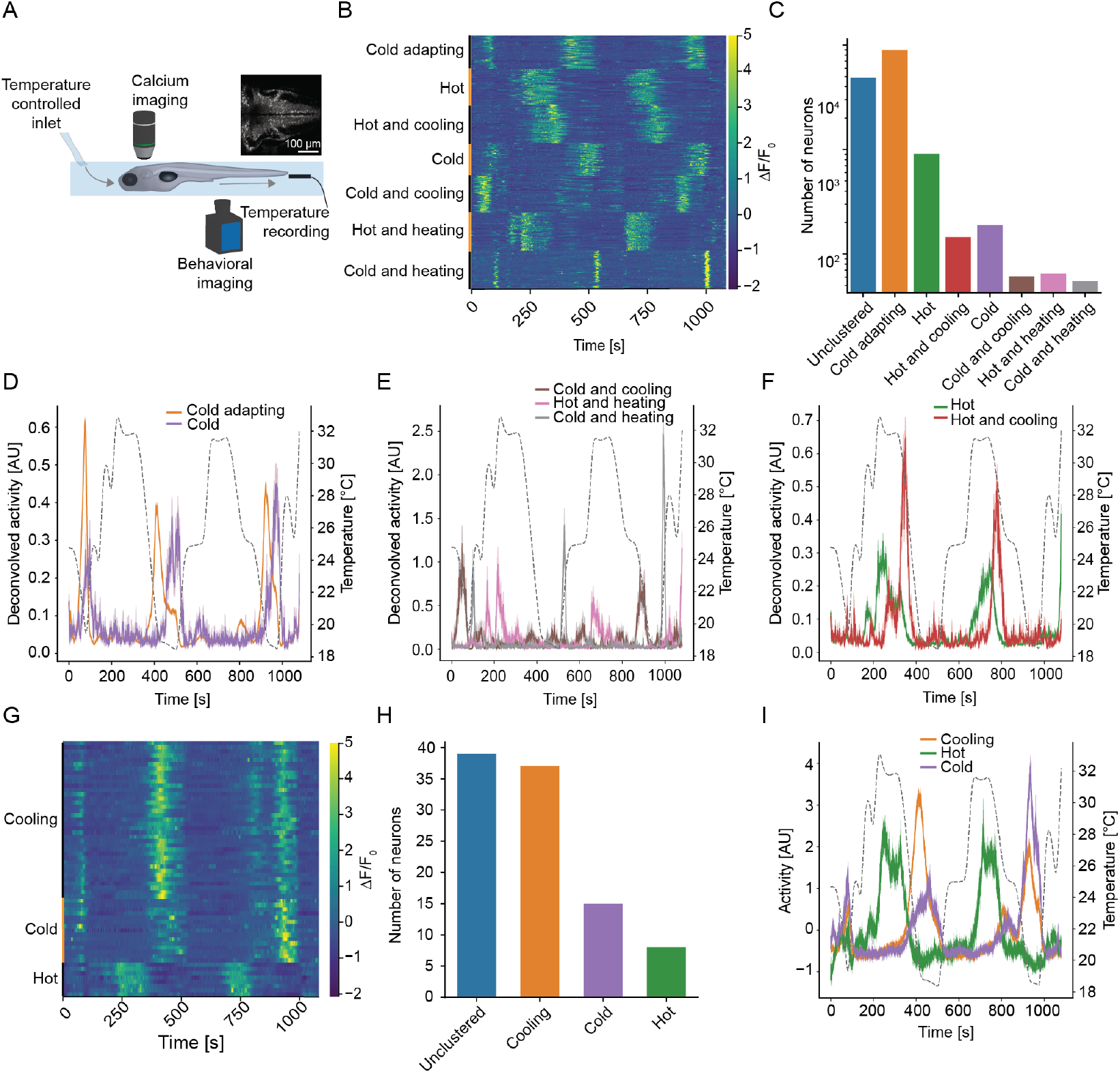
Calcium imaging reveals stimulus segregation across seven response types. **A** Schematic of experimental setup for functional imaging of neurons in the medulla and trigeminal using a custom flow-based setup to temperature-control head-fixed larval zebrafish. **B** Calcium activity of example neurons in identified clusters of temperature responsive neurons in the medulla, clustered according to response correlation. **C** Number of neurons in each identified cluster in the medulla across all experiments. **D-F** Clustered averages of calcium activity of each cluster in the medulla, with the presented temperature stimulus (dotted black line). **G** Same as (B) for neurons in the Trigeminal ganglion. **H** Number of neurons identified in each cluster in the trigeminal ganglion. **I** Clustered averages of calcium activity of each cluster in the trigeminal ganglion, with the presented temperature stimulus (dotted black line). All shaded error regions are bootstrap standard errors across neurons.

After segmenting and deconvolving the imaging data using suite2P ^35^ we used MINE (Model identification of neural encoding ^36^) to identify neurons whose activity could be predicted from the temperature stimulus. Correlation-based clustering ^37^ led to the identification of seven response types (Figure 6B-C). A sizable fraction of neurons remained unclustered (Figure 6C) as these had overall low pairwise correlations (Figure S5B). This was likely because we opted for a lenient cutoff while fitting with MINE to capture as much response diversity as possible while subsequently using the clustering to separate out prominent response types. Each response type encoded distinct features of the temperature stimulus, such as only responding during cold temperatures (Figure 6D) or hot temperatures (Figure 6F), responding to temperature increases while cold (Figure 6E) or responding to temperature increases while hot (Figure 6E). Together with overall low correlations across response types (Figure S5C) this suggested a tiling of stimulus space that could aid in implementing the behavioral algorithm which relied on the identification of improving and worsening contexts. However, no neuron type responded specifically within the preferred temperature range or encoded context across the entire thermal range. Intriguingly, the most abundant neuron type encoded a mixture of cold and cooling stimuli (Figure 6B and D). This overrepresentation of cold stimuli was reminiscent of temperature encoding in the mouse spinal cord in which more neurons encode mild cold than mild warmth ^38^. Further analysis using MINE revealed that all neuron types responded highly nonlinearly. This complexity was largely masked in our previous imaging studies focusing on warm temperatures ^36^, and was a consequence of the very strong rectification present in all response types which either only responded in the warm or in the cold regime.

Previous work suggested that neural temperature responses within the medulla are computed de novo from trigeminal thermosensory inputs^25^. To better understand the relationship between thermosensory responses and the representation of temperature within the medulla, we imaged neural activity within the trigeminal ganglia of larval zebrafish under the same stimulus conditions. Employing the same analysis approach revealed three trigeminal thermosensory neuron types (Figure 6G-H and S5D-E). The most abundant type encoded a mixture of cooling and cold stimuli while the other two types responded to temperatures below 23 °C (cold) or temperatures above 20 °C (hot) (Figure 6I). As in the medulla there was a clear overrepresentation of neurons responding to cold/cooling and the corresponding medullary responses may be directly fed by these thermosensory neurons. As described previously ^25^ responses across types were more similar in the trigeminal ganglion as evidenced by larger correlations and anti-correlations (Figure S5E) than in the medulla (Figure S5C). The decorrelation of neural encoding in the medulla likely serves the purpose of representing important stimulus features involved in behavioral control such as improving and worsening contexts.

In summary, we discovered that larval zebrafish encode temperatures across the entire thermal range. While direct thermosensory representations in the trigeminal were limited, neurons within the medulla were split according to important stimulus features such as context.

### Medullary response classes form a place-like code for thermal gradient navigation

After identification of brainstem temperature representations, we wanted to see how these seven response types could control thermoregulatory behavior (i.e., the avoidance of both hot and cold temperatures). We used MINE to predict the activity of each response cluster using a short sensory history of 2 seconds. On test data, each of these predictions had high predictive power with correlations of at least 0.8 in all cases, indicating that the MINE models generalized well to new inputs (Figure S6A). We subsequently used these models to predict the activity of each response type during the gradient behavior experiments by taking each zebrafish’s experienced temperature as inputs to the MINE models which computed the predicted neural activity (see Figure S6B-I for examples). Analyzing the predicted activity based on the gradient direction and position of swim trajectories revealed a remarkable segregation. The seven response types formed a place-like code within the temperature gradient where the combination of activities marked specific positions and swim directions, such as restricted activity during trajectories in the cold or specific responses on trajectories that started in warm temperatures and moved the fish towards the preference (Figure 7A). This was also apparent when analyzing neural responses according to gradient temperature and movement direction which showed a clear segregation across response types (Figure S6J-P). To test whether activity across the response types could indeed allow zebrafish to infer position and direction within the gradient, we fit linear regression models predicting temperature (position) and temperature change (direction/context) on 80 % of the data and tested it on the remaining 20 %. A success of linear models would indicate that information about position and direction is readily decodable from the seven response types. Across the temperature range, the true temperature could be accurately predicted from neural activity since the slope of the relationship between predicted and real temperatures was close to one (Figure 7B). Overall, prediction errors were smaller for warm than for cold temperatures. However, prediction of temperature change (i.e., gradient direction) was considerably better in the cold (Figure 7C). Here the model revealed that prediction of cooling in the cold had the highest accuracy (Figure 7C top panel). Even small rates of temperature decreases led to predicted temperature changes that were significantly smaller than zero. Interestingly, in the hot regime, heating stimuli were not as well predicted as cooling stimuli in the cold, requiring a larger rate of change to be predicted as significantly larger than zero (Figure 7C bottom panel). However, since larval zebrafish swam larger distances in the hot regime, this would still allow clear identification of the worsening context. Overall, the encoding appeared to favor identification of the worsening context in the cold, which could underlie the high efficiency of avoiding cold temperatures through a strong modulation of reversal modes depending on swim direction.

**Figure 7.**
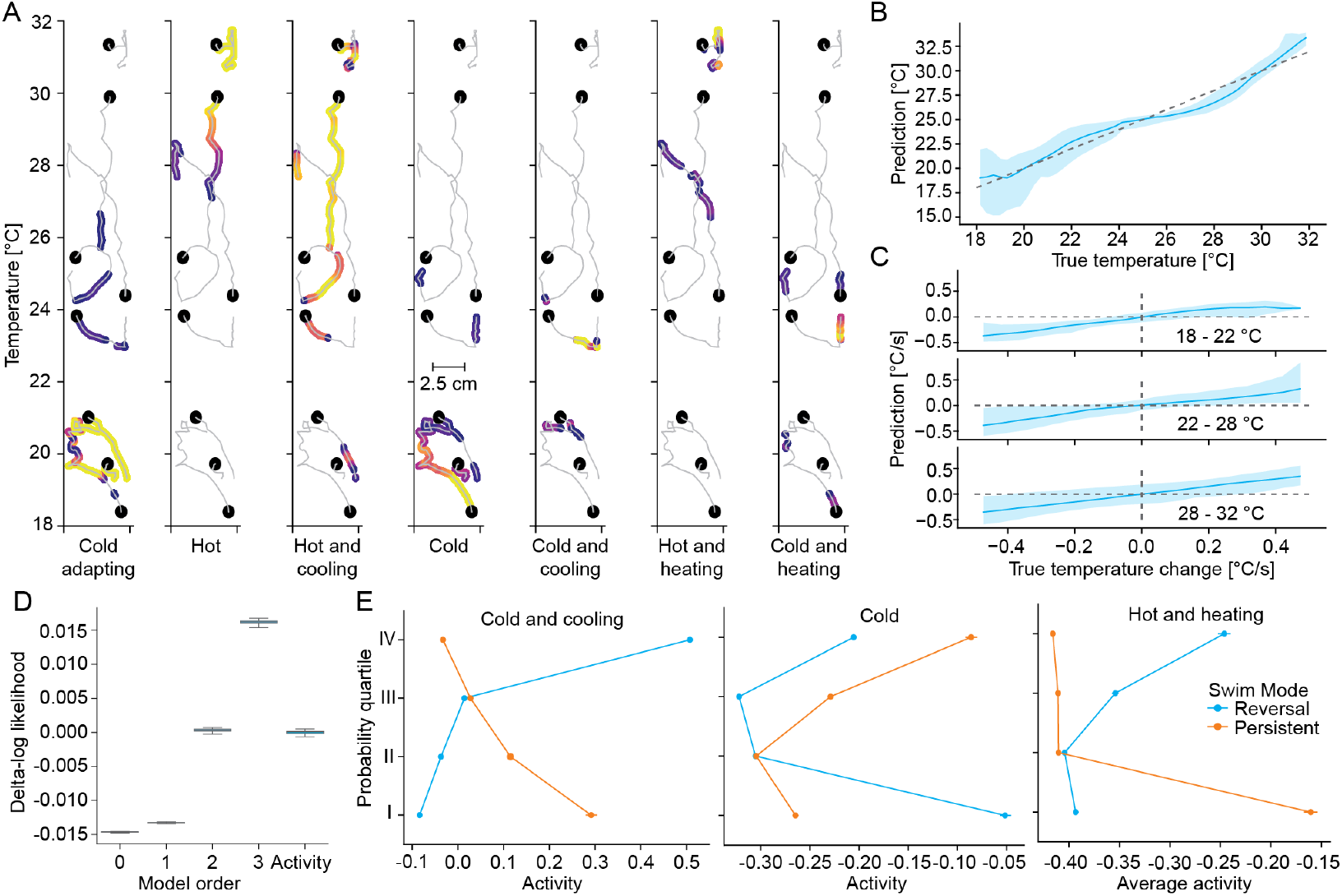
Predicted activity of medullary response types uncovers their role in encoding position and direction within the gradient. **A** For each of the seven response types activity is thresholded along the same set of example trajectories (grey, start of trajectory marked by black dot). Above threshold activity is color coded from 0.5 standard deviations above threshold (dark purple) to 3 standard deviations above threshold (yellow). **B** Comparison of actual temperature of fish with the gradient temperature predicted from a linear combination of predicted neural activity of medullary response types. **C** Comparison of actual change in temperature of fish with the predicted temperature change from a linear combination of predicted neural activity of medullary response types at different temperatures within the gradient. **D** Comparison of goodness of fit of models of different orders and an activity-based model, using the difference in log-likelihood of model predictions on test-data relative to the average of the 2nd order transition model. **E** Influence of activity of neuron type on steady state probabilities of the reversal (blue) and persistent mode (orange). All shaded error regions are bootstrap standard errors across experiments.

To understand whether the neural encoding could control longer-term behavioral features of thermoregulation, we refit the transition model. However, instead of using temperature, temperature change and their interactions as inputs, we designed the model to linearly depend on the activity across the seven response types. This transition model, linear (1st order) in activity space, performed on par with the nonlinear 2nd order transition model in stimulus space in terms of predicting held-out test data (Figure 7D). This indicated that the hindbrain transformed thermosensory stimuli into a code that can be used linearly by downstream circuits to control swim mode transitions underlying gradient navigation, especially cold avoidance. This is in line with the idea that information of importance is represented in a manner that allows linear readout by downstream circuits ^39^–^41^. The transition model made clear predictions about the influence of the activity of individual response types on the steady-state probability of swim modes. Neurons that preferentially mark worsening contexts such as “Cold and Cooling” or “Hot and Heating” responses reduced Persistent mode probabilities while increasing Reversal mode probabilities when active (Figure 7E). Similarly, the Persistent mode was enhanced when neurons marking cold temperatures were active, as predicted from the preferential alignment to the gradient direction in the cold (Figure 7E).

In summary, key behavioral features could be linearly decoded from medullary response types. This included swim-by-swim features such as position and direction/context, as well as longer-term swim modes which are critical for cold avoidance. This strongly suggested that medullary circuits encode the information necessary to control the behavioral algorithm of thermoregulation.

## Discussion

Behavioral thermoregulation is critical for survival across species and evolutionarily ancient. However, despite this fundamental importance, we know very little about the neural and behavioral algorithms that underlie this process in vertebrates. Here we set out to understand thermoregulation in the larval zebrafish by developing a quantitative model of behavior and testing its relationship with neural processing. We used a thermal gradient navigation paradigm to compare the avoidance of temperatures above the preference (hot avoidance) to the avoidance of temperatures below the preference (cold avoidance). This revealed clear differences in strategy. In hot temperatures, larval zebrafish increased overall movement vigor. Paired with increased turning during worsening context this simple strategy was nearly sufficient to explain hot avoidance. In the cold on the other hand, larval zebrafish tended to persistently align to the gradient. A Navigation Model, combining a stimulus-driven Markov model with stimulus driven emission models for swim kinematics, captured these longer timescale behaviors and revealed that nonlinear control of persistent swims along the gradient with intermittent direction reversals was crucial for cold avoidance. This strategy allowed fish to efficiently avoid cold temperatures despite an overall reduction in movement. To assess the neural basis of these behaviors, we performed functional calcium imaging. This revealed seven distinct temperature response types within the medulla of larval zebrafish. These types tiled temperature space, while positions and directions within the gradient could be decoded from their activity. Furthermore, a linear combination of the neural response types could drive swim mode transitions within the Navigation Model. In other words, neural activity explained both instantaneous swim decisions which are based on current temperature and gradient direction as well as context-dependent longer-term strategies which are especially important in the cold.

### Attraction versus avoidance

The goal of behavioral thermoregulation is to seek out a preferred temperature. In theory, one could distinguish two approaches for reaching a preference: active attraction to the preference or avoidance of any temperatures above and below the preference. Since these are two sides of the same coin, it is difficult to disambiguate the two possibilities. However, parsimony speaks to an avoidance-based mechanism of behavioral thermoregulation. At any moment in time, the fish can, at best, know how much warmer or colder it is than the preference and whether it was moving towards (improving context) or away from the preference (worsening context). The spatial location of the preferred temperature cannot be accurately inferred from the gradient without knowledge about the slope and linearity of the gradient. It is therefore likely that thermoregulatory behavior uses local information about the thermal gradient. This local information may then be combined with visual information about landmarks to effectively avoid both hot and cold temperatures.

### Cold avoidance as gradient descent

Larval zebrafish are ectotherm animals. Their nervous and muscular systems are therefore largely at equilibrium with environmental temperature. This creates specific challenges for avoiding cold temperatures ^42^–^44^. As muscle physiology and force generation are impaired, less resources can be mounted in the cold to escape. Larval zebrafish circumvented this problem by using a more deliberate strategy to escape cold compared to hot temperatures. They aligned to the gradient while temperatures were increasing in the cold, thereby picking the most efficient route to their preference akin to a gradient descent algorithm for learning. When encountering decreasing temperatures in the cold however, they initiated direction reversals (Figure 3). These appeared to be a fixed behavioral mode regardless of temperature in which a small number of swims was used to reverse the heading, correcting course to face towards the preferred temperature. These reversals likely corresponded to “U-maneuvers” described as continued turning in the same direction during hot avoidance ^28^. However, the current results suggest that these reversal maneuvers were more critical for cold than hot avoidance. Overall, keeping a persistent direction up the gradient in improving context paired with reversals during worsening context allowed larval zebrafish to avoid cold temperatures despite longer rest times between swims and overall reduced per-swim displacement. This strategy therefore compensated for possible effects of cold on muscle physiology. Endotherms are less impacted by cold temperatures as they are able to raise their body temperature above environmental temperatures. However, thermal responses in mammals overrepresent cooling stimuli ^38^ and humans are more sensitive to cooling than heating stimuli ^45^–^47^. This parallels our behavioral and neural findings in zebrafish, where cold and cooling responses were highly overrepresented in the medulla. This suggests evolutionary conservation of the importance of cold responses and cold avoidance from ectotherms to endotherms.

This overrepresentation of cold stimuli was already present in the trigeminal ganglia, in which two neuron classes encoded temperatures below 23 °C (Figure 6). However, how zebrafish detect these temperatures is unclear. Teleost fish lack the canonical receptor for ambient cold, TrpM8^48,49^ and it has been speculated that fish might not actively avoid cold temperatures since most species can tolerate cold^50^. However, cold avoidance was clearly present in zebrafish (Figure 1) despite the ability to survive at temperatures as low as 14 °C^51^. The ability to navigate away from cold temperatures is clearly advantageous for tropical fish that live in environments with fluctuating temperatures such as zebrafish ^52^ to keep their metabolism functioning optimally. Given the presence of cold activated thermosensory neurons, it would be of great interest to identify the corresponding cold receptor to gain a clearer understanding of the evolution of thermosensation.

### Instantaneous modulation and behavioral modes

Larval zebrafish thermoregulation combines elements of a biased random walk (changes in swim kinematics with temperature and temperature change and increases of reversals) with a more deliberate strategy of aligning to the gradient direction in the cold. The most simplistic, yet inefficient, strategy for seeking out a preference is orthokinesis or diffusional trapping. It relies on increasing movement with increasing distance from the preference, while increasing rest at the preference. This strategy exclusively aids hot avoidance in larval zebrafish ^26^ and was observed when isolating the effect of constant temperature on swim selection (Figure 2A-C). A more efficient strategy, employed in bacterial chemotaxis and thermotaxis is a biased random walk^53–55^. Here, unfavorable changes (worsening context) lead to increases in reorientation (turn angles in the case of zebrafish). This strategy, together with orthokinesis, is largely sufficient in explaining the effective hot avoidance larval zebrafish display^23^,28 (Figure 2D-F). Larval zebrafish additionally repeat the same turn direction during worsening context, enacting near ballistic reversals or U-maneuvers ^28^ which enhances the effectiveness of reorientations. The ability to reverse direction across multiple turns enhanced hot avoidance and was critical for cold avoidance which in addition relied on the fish persistently aligning to the gradient direction (Figures 3-5). How these longer-term strategies are controlled is still unclear, however potential neural substrates have been identified in the zebrafish. A putative oscillator within the medulla underlies the maintenance of turn directions during exploratory motion and phototaxis ^33^,56,57 while habenular circuits have been implicated in persistent turning during hot avoidance ^28^.

### The neural basis of thermoregulatory decisions

As in other vertebrates, behavioral thermoregulation in zebrafish relies on a distributed network of temperature encoding neurons across various brain regions ^22^. Specifically in larval zebrafish, trigeminal thermosensation, as well as computations within medullary circuits and the preoptic area and habenula have been implicated in hot avoidance ^25^,28,36,58. However, neural responses to temperatures below the preference have not been previously recorded or analyzed. Here, we used a new experimental design to perform functional calcium imaging across the same thermal range (18-32 °C) in which we tested zebrafish gradient navigation. We focused our imaging efforts on the medulla since circuits within this brain region control thermosensory motor transformations during hot avoidance ^25^. Neurons within the medulla encoded temperature across seven functional types (Figure 6). All of these types were preferentially active at either warm or cold temperatures with a clear zone of overlap. One cell type each was activated by warm or cold temperatures while the other five types preferentially responded to temperature change, three in the cold, two in the warmth. The most abundant response type encoded a mixture of cold and cooling signals (cold adapting neurons). We used a convolutional neural network method we recently developed (Model Identification of Neural Encoding, MINE, ^36^) to generate predictive models that allowed us to infer the response of each neuron type based on temperature experienced during zebrafish gradient navigation. This allowed us to analyze the contribution of these neuron types to hot and cold avoidance during behavioral thermoregulation (Figure 7). Intriguingly, the seven response types formed a place-like code. A linear model of the activity could inform fish about position and direction within the gradient. In the cold, rates of cooling of less than 0.2 °C/s were readily decoded from activity. Temperature increases on the other hand were only decodable at rates larger 0.3 °C/s, independent of absolute temperature. This is reminiscent of increased sensitivity to cooling stimuli in mammals ^38,45^ and likely underlies the efficient behavioral strategy of cold avoidance. Temperature representation across the seven response types also formed an efficient code for guiding swim mode transitions, a crucial component of cold avoidance. Reversal and persistent swim modes were especially influenced by neurons encoding worsening context. Namely, reversal probabilities were increased when these neurons were active while persistent swimming in the gradient direction was suppressed. It is therefore conceivable that these hindbrain neurons control both moment-to-moment modulations in swim kinematics as well as longer term behavioral strategies during gradient navigation.

### Outlook

Behavioral thermoregulation is widespread across the animal kingdom ^8^. When given a choice, even endothermic mammals will use behavior to seek out preferred temperatures ^13,16,59^. This behavioral component dampens fluctuations in core temperature which indicates an intricate interplay between behavioral and autonomous regulation of body temperature^16^. Using larval zebrafish, in which behavior is the only output of the thermoregulatory control system, we investigated behavioral and circuit principles underlying thermoregulation. We specifically focused on how larval zebrafish control and enact a universal building block of any thermoregulatory strategy: the detection of worsening and improving conditions and their relationship to reorientations and persistent movement. Notably, worsening vs. improving contexts are dependent on both changes in temperature as well as absolute temperature, since the valence of a temperature change depends on the relation of the current temperature to the preference. E.g., heating stimuli signal worsening context above the preference while they signal improving context below the preference. Here we could show that larval zebrafish solve this integration through a distributed code in which neurons are preferentially active at low or high temperatures, signaling both absolute temperature and temperature change. This place-like code allows the animal to enact movement decisions required for behavioral thermoregulation.

We focused our functional analysis onto the medulla since it is the first integrative center of skin temperature in vertebrates, through structures such as the nucleus of the trigeminal and the lateral parabrachial nucleus (lPB)^9^,^10^. Furthermore, in zebrafish temperature encoding neurons within the medulla are sufficient to explain critical features of hot avoidance ^25^. It is interesting that already at this stage an efficient code arises that can be linearly read out to control various aspects of thermoregulatory behaviors. However, we currently do not distinguish ascending structures within the hindbrain such as the lPB from processed descending outputs that may directly interact with pre-motor neurons within the medial and anterior medulla. These outputs could arise after processing within zebrafish forebrain structures such as the preoptic area (POA) and the habenula. The POA is a major integrative center in mammalian thermoregulation ^10^ and ablation of the POA in zebrafish reduces the enhancement of turn angles during worsening context in hot avoidance ^28^. We previously identified temperature encoding neurons within the zebrafish POA which encode temperature fluctuations on longer timescales^25^,36. As in mammals, the POA may therefore serve a role of setting or modulating the thermal preference in larval zebrafish rather than directly controlling the behavioral strategy. POA outputs may specifically control threshold temperatures at which “warm” and “cold” responsive neurons in the medulla activate. Such modulation would thereby shift the temperature at which activity across these classes separates. This separation is crucial for the linear readout of worsening and improving context and Figure 7B reveals that the relationship between true temperature and temperature decodable from the neurons flattens for temperatures within the zone of activity overlap. An expansion of this zone may lead to the observed reduction in turning by worsening context observed in the POA ablations. This hypothesis could be tested by combining POA ablations with calcium imaging and behavior across the thermal range since effects would likely be asymmetric in the hot and the cold due to differences in the sensitivity of detecting temperature change.

During hot avoidance, the habenula has been implicated in maintaining direction across consecutive turns ^28^ and in mammals habenular neurons have been implicated in cold aversion ^60^. Ablations of neurons within the dorsal part of the zebrafish habenula led to a reduction in the maintenance of turn angles which could directly impact reversals ^28^. If this is a general function of the habenula we would expect that it leads to strong defects in cold avoidance since this behavior is more reliant on long-term behavioral modes than hot avoidance. Another intriguing role of the habenula could be in controlling alignment to the gradient direction. We found that larval zebrafish preferentially align to the gradient direction in the cold, and this is likely what allows them to efficiently swim away from cold temperatures despite a reduction in overall movement vigor. Recently, a heading direction network was discovered in larval zebrafish which involves habenular inputs to the interpeduncular nucleus to control the tracking of landmarks^61,62^. It is interesting to speculate that heating stimuli in the cold (improving context) could influence landmark representation allowing zebrafish to align to the gradient. In fact, weaker alignment in constant-temperature experiments suggests an integration of visual and thermal information in creating the observed gradient alignment. We recently discovered mixed selectivity neurons that jointly encode swim displacement and temperature change^36^. Such neurons could inform larval zebrafish about the slope of the gradient and thereby provide critical inputs to the heading direction network about the direction of steepest ascent, i.e., the gradient direction. Integration with visual landmarks could then allow effective maintenance of optimal trajectories.

### Limitations

Due to technical limitations, we were not able to directly assess the thermal preference of larval zebrafish. This would have required a larger thermal gradient and a larger chamber, as zebrafish started to behave erratically in steep gradients. However, the modulations of kinematics and behavioral modes we observed can be used to estimate the preferred temperature range to be between 23-27 °C. Within this range there is little modulation of turn magnitude and reversal probability by temperature change (Figure 1 and 3). The same is true for the length of persistent trajectories which is consistently modulated by increasing vs. decreasing temperature only above 27 °C and below 23 °C (Figure 3E). Using the Navigation Model to simulate navigation across the entire thermal range while maintaining the overall slope of the gradient, suggested a preference around 25 °C which is in agreement with the observations about behavioral modulation (Figure S4K). On a neural level, the slope between true temperatures and temperatures predicted based on activity flattened between 24-28 °C (Figure 7B) agreeing with the preference range suggested by behavioral features.

The limitations in capturing cold avoidance of the Navigation Model are likely a consequence of treating reversals as a consecutive series of individual turns rather than a compound maneuver as well as the inability to enforce gradient direction alignment within the model. Specifically, by construction a Markov model will assume that the probability of the next state only depends on the current state and not any previous states. Reversal maneuvers are therefore interrupted within the model, even though fish data suggests that reversals generally run to completion in a fixed number of swim bouts. Furthermore, the lack of an input related to gradient direction meant that the persistent state would lead to less efficient swims within the model compared to the real fish. These choices were made deliberately, until we have evidence of behavioral and sensory mechanisms that could underlie compound reversals and gradient alignment respectively.

## Methods

Animal handling and experimental procedures were approved by the Ohio State University Institutional Animal Care and Use Committee (IACUC Protocol #: 2019A00000137-R1).

### Code and data availability

All code used in this study is available in the following repositories:

Analysis of zebrafish behavior: https://github.com/haesemeyer/thermonavi_behavior

Behavior and model-based simulations: https://github.com/kaarthik-balakrishnan/thermonavigation-simulation

Fitting of Navigation model using STAN: https://github.com/haesemeyer/thermonavi_stan Analysis of calcium imaging data: https://github.com/haesemeyer/thermonavi_imaging Raw experimental data has been deposited to DANDI in NWB format:

- Cold regime gradient experiments https://dandiarchive.org/dandiset/000707
- Hot regime gradient experiments https://dandiarchive.org/dandiset/000708
- Constant temperature behavior 16 °C https://dandiarchive.org/dandiset/000697
- Constant temperature behavior 18 °C https://dandiarchive.org/dandiset/000698
- Constant temperature behavior 20 °C https://dandiarchive.org/dandiset/000699
- Constant temperature behavior 22 °C https://dandiarchive.org/dandiset/000700
- Constant temperature behavior 24 °C https://dandiarchive.org/dandiset/000701
- Constant temperature behavior 26 °C https://dandiarchive.org/dandiset/000702
- Constant temperature behavior 28 °C https://dandiarchive.org/dandiset/000703
- Constant temperature behavior 30 °C https://dandiarchive.org/dandiset/000704
- Constant temperature behavior 32 °C https://dandiarchive.org/dandiset/000705
- Constant temperature behavior 34 °C https://dandiarchive.org/dandiset/000706
- Trigeminal calcium imaging https://dandiarchive.org/dandiset/001339
- Medulla calcium imaging https://dandiarchive.org/dandiset/001337

Processed data has been deposited on Zenodo:

- Processed gradient behavior data including medulla response predictions https://doi.org/10.5281/zenodo.14902233
- Processed constant temperature behavior data https://doi.org/10.5281/zenodo.14902292
- Model simulation data https://doi.org/10.5281/zenodo.15028245
- STAN posterior samples of navigation model parameters https://doi.org/10.5281/zenodo.14902972
- MINE weights of fits of medulla response types https://doi.org/10.5281/zenodo.15007957

### Fish strains

All experiments were performed in unpigmented offspring of incrosses between mitfa +/-; Elavl3-H2B:GCaMP6s ^34^ animals at 5-7 days post fertilization.

### Gradient setup and experimental protocol

The gradient setup consisted of an aluminium chamber which was machined with two swim-lanes and two temperature measurement-lanes in the center. Each swim lane measured 200 mm in length and 50 mm in width with a depth of 5 mm. The measurement lanes had the same length and depth but a width of 5 mm. The base of the chamber was fitted with 6 Peltier elements (Custom Thermoelectric, USA), one per swim-lane at the front, middle and back of the chamber along the long edge. Each pair of these Peltier elements (front, middle, back) was independently controlled by a custom written software PID controller (C# Microsoft, USA) that compared the current temperature in the measurement lane to the desired set temperature. The Peltier drive current was set by the PID controller through a motor driver (Pololu, USA) via analog outputs from a National Instruments PCIe 6343 DAQ board (National Instruments, USA). The current temperature was measured along the temperature-lanes using thermistors (Littlefuse USP16673, Digikey, USA) that were placed at the centers of the respective Peltier elements. The swim-lanes were filled with E3 medium and the temperature-lanes were filled with distilled water to protect the thermistors. The waste heat generated by the Peltier elements was dissipated using a custom-machined copper block, followed by standard CPU cooling fans. The temperature was set using a custom written UI that displayed the current temperature and the set temperature. Temperature measurements were also continuously recorded to disk by the same program. The set temperature was typically reached quickly and gradients stabilized within 30-60 minutes, beyond which there was little fluctuation from the set temperature (*< ±*0.5 °C). Experiments were only started after the gradient or constant temperature profile had stabilized.

The gradient chamber was illuminated from the sides by two white light panels (Nanlite, USA) situated on either side of the chamber. A Blackfly S BFS-U3-28S5M camera (FLIR, USA) was placed above the chamber taking images at a framerate of 100 Hz with a resolution of 0.14 pixels/mm.

For each experiment, a single larval zebrafish was introduced into each swimlane and images were acquired with custom written software (C# Microsoft, United States) and custom tracking algorithms were used to find the position and heading direction of the fish as previously described ^24^ over the course of a 30-minute experiment. Due to practical limitations in data storage, tracking was performed online and raw camera images were not stored to disk. All further analysis was performed on the stored positions and heading directions of the larvae.

### Imaging setup and experimental protocol

Functional calcium imaging was performed on a custom-built two-photon microscope ^36^ at 2.5 Hz with a pixel resolution of 0.78 *µ*m*/*pixel (medulla/hindbrain) or 0.33 *µ*m*/*pixel (trigeminal ganglion) and at an average power of 12 mW at sample at 950 nm (measured optical resolution of the system is *<* 1 *µ*m lateral and *<* 4 *µ*m axial). Imaging was either in the medulla or the trigeminal ganglion. Larval zebrafish were mounted in a custom designed chamber in which water was flowing through a channel. To generate different flow temperatures, an inline flow heater (Warner Instruments, USA) with associated controller was used. Control temperatures were calibrated such that at sample temperatures in the range from 18 °C to 32 °C were reached. Temperatures at sample were continuously monitored through a thermistor (Warner Instruments, USA) placed behind the larva and were recorded during the experiment. During imaging scan stabilization along the z-axis was used to avoid artefacts induced with heating and cooling the agarose with the flowing water.

For temperature stimulation, within each imaging plane a 18 minutes long stimulus was presented to the larval zebrafish. The stimulus was fixed across all imaging planes and consisted of sine wave fluctuations, ramps and temperature holds. Imaging planes were spaced 5 *µ*m apart and 10-20 planes were imaged in each fish for experiment in the medulla and 4-6 planes for trigeminal imaging experiments.

### Data analysis and modeling

#### Swim bout identification and kinematics

Fish position traces in x and y were filtered using a gaussian filter with a standard deviation of 2 frames (20 ms) and the instantaneous speed was subsequently calculated as the per-frame euclidean distance traveled. The instantaneous speed was used to identify bouts via peak detection. Peaks with a minimum speed of 5 mm/s and a minimum width of 50 ms were identified. Bout starts and ends were defined as the first frame before/after the peak where the acceleration/deceleration dropped below 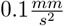. According to these bout calls, kinematics were defined. The interbout-interval (IBI) was defined as the time between the end of the previous bout and start of the current bout. The turn angle was defined as the change in angle between the average heading angle of five frames following the bout and five frames preceding the bout, such that all turns were assumed to be smaller than 180 degrees. Similarly, the bout displacement was defined as the euclidean distance between the position after the bout and prior to the bout.

For turn magnitude and direction correlation analyses, turns were separated from straight swims using a cutoff of *±*5 ^*°*^. This cutoff was chosen as it was the average standard deviation of the straight swim gaussian across all swim modes.

#### Exclusion of swims near the edge of the arena

Since the edges of the arena likely influence swim kinematics, e.g., by preventing turning into a wall, all swim bouts that were within 4 mm (one fish length) of either of the four sides of the chamber were excluded from all further analysis.

#### Definition of continuous trajectories

For all correlation based analysis and Markov Modeling, only continuous swim trajectories were considered. Continuous trajectories were defined as a collection of successive bouts with inter-bout intervals below 2 s. We note that some trajectories at colder temperatures were likely cut because of this interbout interval rule even though they were continuous. However, this stringent cutoff was chosen to ensure that correlations and Markov transitions were only evaluated across a continuous block of time and not influenced by the tracker intermittently losing the fish. Only trajectories with at least three swim-bouts were used for Navigation model fitting and testing.

#### Assignment of stimulus features to swim bouts

For stimulus based analysis, model fitting and simulations, two stimulus variables were assigned to each bout: The temperature at the start of the bout as well as the temperature change experienced over the previous bout. This value, rather than average temperature change over a fixed time interval, was chosen, since swims are the only time where fish will experience changes in temperature within a gradient.

The temperature change across the previous bout was used to define “heating” and “cooling” directions for single bout analysis (Figure 1). Specifically, the median absolute temperature change (*∽* 0.52 °C*/*s) was used as a cutoff, such that bouts with a previous temperature above this threshold or a negative temperature change below the negated threshold were considered to have followed a “heating” or “cooling” bout respectively. For trajectory based analysis (Figure 3) “heating” and “cooling” were defined based on the movement direction being either towards the hot or the cold end respectively. The different thresholding was chosen for the two analyses since one was concerned with the influence of the stimulus on kinematics which likely depended on the stimulus crossing a threshold while the latter was concerned with how fish change direction within the gradient based on direction of travel.

#### Defining maneuvers and swim modes

To analyze longer-term behavioral strategies as well as behavior in relation to the gradient direction, additional features of swim bouts were defined based on the displacement direction of swims within the chamber rather than the heading angle of the fish. Specifically, for each swim displacement the “gradient direction” was defined as the cosine of the angle of the displacement relative to the gradient direction, such that swims in the heating direction had a negative gradient direction while swims in the cooling direction had a positive gradient direction (since increasing y-coordinates were associated with colder temperatures within the chamber). The absolute value of gradient direction (absolute cosine of the angle) was defined as the “gradient alignment” which is 1 for any swim displacement directly along the gradient direction and 0 for any swim perpendicular to it. Based on these metrics, swim displacements within a *±*45 ^*°*^ cone around the gradient direction (absolute cosine *>* 0.71) were defined as aligned swims.

Based on this labeling swim modes were defined in the following manner. A “reversal” trajectory was defined as the set of all bouts that started with the fish aligned in either the positive or negative direction followed by a set of bouts that ended with a displacement in the opposite aligned direction. 90 % of such reversals were completed within 10 swim bouts and 10 swim bouts was therefore chosen as the maximum length to call this trajectory a “reversal maneuver.” A “persistent” trajectory was defined as the set of all bouts that were consistently aligned to the same gradient direction with a minimum length of two swim bouts. All remaining swim bouts were assigned as “general bouts.”

By construction, every bout was classified into one of these categories. These categories were therefore used to define the current swim mode as either reversal, persistent or general. Those labels were used to calculate reversal initiation probabilities, the average length of persistent trajectories, etc. The swim modes also formed the basis of the Navigation Model.

#### General approach to model fitting and testing

All models were fitted using Hamiltonian Markov-Chain Montecarlo via STAN^31^. For model-fitting the respective design matrices *X* were orthogonalized using QR-Decomposition within STAN. Model parameters *θ* were fit in this space and subsequently put into the original space using the inverse transform. Linear dependencies were thus removed from the fit while still obtaining parameters *β* that could be used on raw data during tests and simulations.

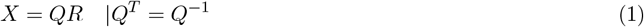

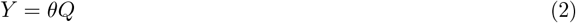

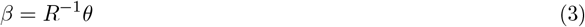

Swim bouts were grouped into trajectories and 80 % of all trajectories were used to fit models (Markov model transitions, emission models) while the remaining 20 % were used for model testing. Trajectories were split into the train/test set by adding 4 consecutive trajectories into the train set, followed by adding one trajectory into the test set and so on. Therefore, each set was made up of trajectories across all fish. Model tests were therefore used to assess generalization of fits w.r.t. an average fish, not across fish. For each model, 4000 draws of parameters were generated from the posterior distribution across four independent chains. For all model tests the individual draws were used (one draw per test across 100 test iterations) rather than testing performance on the parameter average.

#### Markov model of swim modes

The Markov model was defined to capture transitions between the three swim modes, reversal mode (r), persistent mode (p) and general mode (g). The transition probabilities between modes *M* were set by a Generalized Linear Model (GLM) with the current temperature, temperature change across the previous bout and their interactions as inputs such that *p*(*M*_*t−*1_ *→ M*_*t*_) = *f* (*T*, Δ*T*) with the parameters of *f* (*T*, Δ*T*) depending on the model order *k*. Of the three transitions from each swim mode the log probability of one was pinned to zero (to ensure identifiability of the resulting model), leading to the following definition of transition probabilities with the 2nd order model as an example:

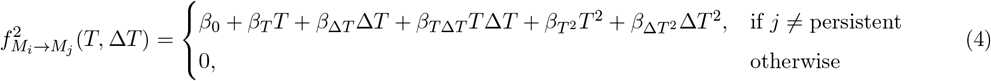

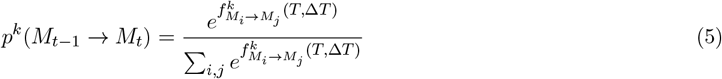

Steady state probabilities *p*(*M*) for each mode *M* for different inputs were computed by exponentiating the transition matrix Θ 100 times. This approach was chosen for efficiency over repeated eigen-decomposition, however we note that the steady state probabilities can be computed analytically as the right eigenvector with the highest eigenvalue (1) of the transition matrix. These approaches are equivalent as long as the transition matrix does not describe an oscillatory system.

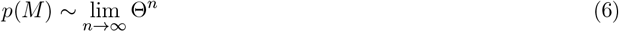

For model testing, a “background” or chance model was constructed to evaluate the importance of GLM controlled transitions on predicting swim modes. The overall probabilities of each swim mode across the training data were used in this model to predict the current state while the Markov model was used in the main model test to predict the next state given the current state and the transition matrix computed from the GLM of the chosen order. The log-likelihoods of the test data given the chance model *ll*_*chance*_ and of the transition model of order 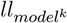 were then computed as follows:

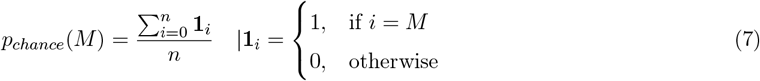

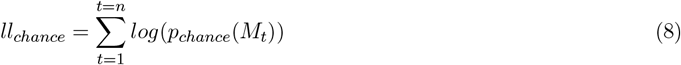

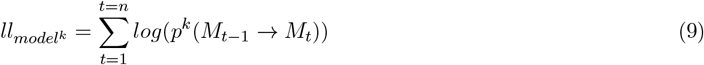

Importantly, these quantities were only computed across transitions within trajectories, not across trajectories. In other words, for both *ll*_*chance*_ and *llmodelk* computations were limited to elements two to n in each individual trajectory and subsequently summed across trajectories.

#### Emission models of swim bout kinematics

To capture swim bout generation dependent on swim mode as well as the stimulus and behavioral history the following emission models were fit. For each model, the training data was first split according to swim mode and subsequently separate emission models were fit for each mode. The rationale for this approach was that different modes were likely generated by eliciting different swim bouts. At the same time the expectation was that within each swim mode, bout kinematics would be influenced by the stimulus as well. Hence separate GLMs relating stimulus features to kinematic distributions were fit for each mode.

#### Turns

Turns were modeled as a Gaussian-Gamma mixture model similar to Dommanget-Kott^27^. Straight swims were captured using a Gaussian distribution centered at 0 with a standard deviation *σ*_*s*_ fit on the data. Left and right turns were modeled as mirror-symmetric Gamma distributions with parameters shape *α*_*ϕ*_ and rate *β*_*ϕ*_ fit on the data. *σ*_*s*_, *α*_*ϕ*_, *β*_*ϕ*_ were fixed for all turns within each mode but were fit independently for each mode. Within each mode, turn distributions were modeled as mixtures of these distributions with the mixing coefficients Θ dependent on the current temperature *T*, temperature change Δ*T* of the previous bout as well as the previous turn angle 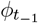, according to:

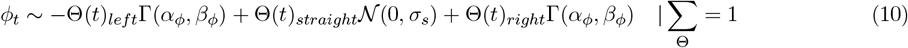

with

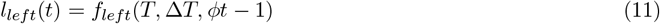

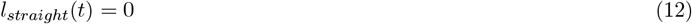

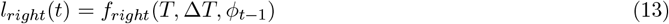

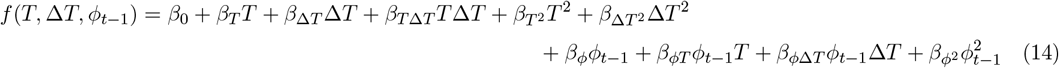

and

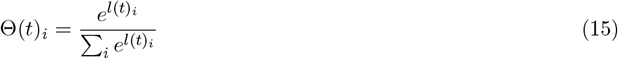

#### Displacements

Displacements *d* were modeled as a single Gamma distribution since displacements are strictly positive. While not a perfect description of displacements out of the standard distributions, displacements were best fit with this distribution. The distribution was parameterized using rate *β* and average *µ* such that:

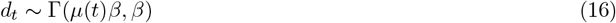

This parameterization was chosen under the assumption that larval zebrafish control the average displacement. It also yielded more stable fits than a standard parameterization according to shape and rate. The rate parameter *β* was fixed for each swim mode while *µ*(*t*) was modeled as a function of the current temperature *T*, temperature change Δ*T* of the previous bout as well as the previous displacement *d*_*t−*1_, according to:

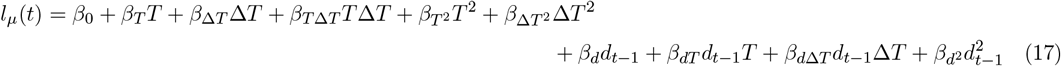

and

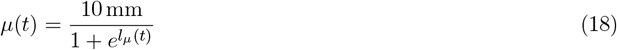

In this formulation 10 mm was chosen as the maximal average displacement. Given that average displacements irrespective of the stimulus never exceeded 4 mm this was a mild constraint on the fit.

#### Inter-bout intervals

Interbout intervals were well described as Gamma distributions and fit in the same way as bout displacements with the only difference being that the average bout displacement was modeled as a function of temperature and temperature change only, since auto correlations were low for inter-bout intervals. Therefore:

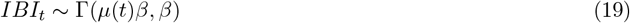

with

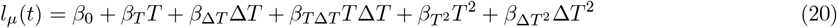

and

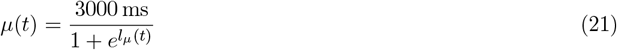

Model tests for the emission models were performed in the following manner. For each mode a background/chance model was constructed in which the appropriate distribution (Gamma or Gamma/Gaussian mixture) was fit on all training data within the mode. The likelihood of this chance model was computed on the test data by first conditioning on the mode and then computing the likelihood of the observed kinematics according to this fixed model. Comparison to tests of the GLM controlled kinematic emission models therefore tested the improvement in log-likelihood given the stimulus control of the kinematic emission model.

#### Simulations

Based on experimentally identified swim bouts and bout parameters along with their dependence on temperatures, the fish was simulated as a virtual particle moving in a gradient. Since all analysis of the fish in experiments assumed the fish as a point particle with a defined centroid and heading angle, the virtual fish in simulations was modeled as a point particle with an associated vector as a heading direction as well. A virtual chamber was defined according to the exact same dimensions as the experimental setup, with 1464 pixels along the length and 320 pixels along the width. The spatial resolution was defined as 7 pixels/mm, identical to the image resolution during experiments. The particle was started at a random point in this chamber with a random heading direction, and was moved along the chamber by picking “bouts” from either the actual experimental distribution (constant temperature and temperature+temperature change simulations) or generated bouts from the Markov gradient navigation model. Each simulation was run for a length of 180,000 frames at 100 Hz, the same length as the 30-minute experiments. Two kinds of gradients, similar to the experimental setup were imposed on the chamber, one ranging from 18 °C at the rear end (y=1464) to 26 °C in the front end (y=0), to another set ranging from 24 °C at the rear end and 32 °C at the front end. The slope of both of these gradients was therefore the same as the slope of the experimental gradients 0.4 °C/ cm. Additionally, a “full” gradient was simulated, ranging from a temperature of 18 °C to 32 °C. The length of the chamber was increased to 2562 to match the slope of the gradient to the experiments and other simulations.

For the constant temperature simulations, swim bouts were selected as follows. The current temperature of the particle within the simulation was used to pick a swim-bout from a corresponding constant temperature experiment. As these experiments were run only in 2 °C intervals, bouts were probabilistically picked from the two closest available experimental conditions, such that the ratio of the probabilities depended on the ratio of the distances to the three nearest temperatures *a, b* and *c*:

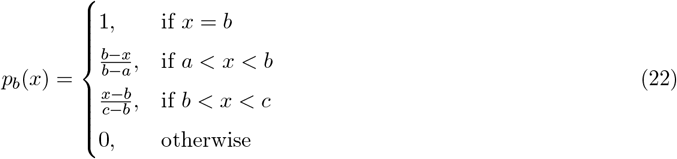

Once the temperature bin was determined, a random turn angle, bout displacement and interbout interval were picked from the corresponding constant temperature experiment. Using these quantities, the particle was moved to a new position in the chamber, first updating the heading angle, followed by a displacement along the new direction, followed by an interbout interval. Edge conditions were resolved such that whenever a movement would take the particle across the edge of the virtual arena, its direction of movement was aligned with the edge. This approach was chosen to mimic edge tracking behavior in larval zebrafish.

For the simulations combining temperature and temperature change, bout data from the gradient experiments was binned according to a) the temperature at the start of the bout and b) the temperature change experienced during the preceding bout. This bin width for temperature was chosen to be 0.5 °C and the bin width for temperature change was chosen to be 0.05 °C. The virtual particle was again started at a random position in the chamber with a random heading angle and a previous temperature change of 0 °C. Since the bin sizes were small, the bin with the closest matching temperature and temperature change was chosen to instantiate the next bout. The temperature change induced by this bout was subsequently used to find the bin for picking the next bout and so on. Using these quantities, the particle was once again moved to a new position in the chamber with simulations proceeding as in the constant temperature case.

Simulations using the Markov model proceeded in the same manner as the other simulations. The major difference was that swim bout were chosen according to a parametric model instead of from raw data bins. At the start of the simulation the particle was set to be in the General swim mode. Subsequently swim bouts were chosen in the following manner. The Markov model was used to implement a probabilistic state transition with the transition matrix being computed using the current temperature and the temperature change across the previous swim bout. For computation a random parameter sample was drawn from the posterior distribution. After the state transition, a turn angle, displacement and interbout interval were drawn according to the probability distributions generated by the emission models for the newly selected state. Characteristic parameters of the distributions (see: Emission models of swim bout kinematics) were computed using temperature, temperature change, previous bout angle and previous displacement by drawing a random parameter sample from the swim modes posterior distributions. Subsequently a random turn angle, displacement, and interbout interval were drawn from the emission distributions. The angle and displacement were used to implement movement of the particle while the interbout interval was used as a waiting time until the next swim mode transition and bout implementation.

#### Comparison of simulation and experimental gradient outcomes

To measure how well simulations approximated gradient navigation behavior, the Kullback-Leibler (KL) divergence between each fish-average experimental gradient distribution (18-26 °C gradient ant 24-32 °C gradient) and the average across 200 simulations in the same gradient was computed. Since it was not the goal to compare simulation performance for edge-tracking behavior, data within 0.5 °C (1.25 cm) from each edge were excluded from the analysis. The remaining data was divided into bins with a width of 0.14 °C. After calculating the proportion within each bin *b* for the experimental fish data *P* (*b*) and the simulation data *Q*(*b*), the KL-divergence was computed according to:

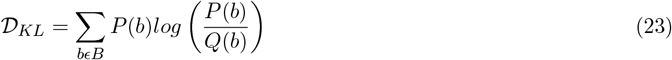

As a comparison, the KL divergence of a uniform distribution to each experimental distribution was computed as well. This would be the distribution obtained when simulating a randomly moving particle.

#### Identification of neuron response types

Raw imaging data was processed using Suite-2P to extract individual neural responses in each plane as well as deconvolve the calcium traces. Suite-2P was fine tuned to properly identify neurons in our imaging setup. Subsequently MINE was used to identify neural responses that could be predicted based on the temperature stimulus ^36^. A correlation cutoff on test data of *r* = 0.6 was used to decide that a neuron’s activity was related to the stimulus. For hindbrain data all neurons passing threshold were used for subsequently clustering analysis. For trigeminal data, since our imaging planes also contained part of the hypothalamus, an anatomical mask was applied before the MINE fitting step to only select responses of trigeminal neurons.

Temperature stimulus related neurons were subsequently clustered using a greedy correlation based approach ^37^, with a correlation cutoff of 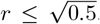. For hindbrain data a minimal cluster size of 25 was imposed and each cluster had to be present in at least 25 independent imaging planes. For trigeminal data, this criterion was relaxed to a minimal size of five neurons, present across at least three imaging planes. Clustering was performed on the raw fluorescence traces. For hindbrain data cluster average activity was reported using the deconvolved data. This approach was chosen to eliminate noise during clustering while improving timing accuracy in the reported responses to aid activity prediction during behavior (see below). This approach was not possible for the trigeminal data. Since the trigeminal ganglion is located very ventrally, imaging data obtained in the ganglion was too noisy to allow for deconvolution.

#### Prediction of neural activity during swimming and activity based modeling

MINE yields models with high predictive power allowing to predict how neurons were responding during the behavioral experiments. To exploit this fact, we trained MINE convolutional neural network (CNN) models to predict the deconvolved activity of each response type in the medulla based on the temperature stimulus. To better capture behaviorally relevant response timescales, only two seconds of stimulus history were used in these predictions. The depth of the CNN was increased to four layers which was required to get predictive power on some of the response types. The models were subsequently used to predict the response of each of the seven response types in the medulla at each timepoint in the gradient experiments by using the temperature experienced by experimental fish exploring the gradient.

To test whether gradient position and direction could be decoded from the activity, linear regression models were trained to predict the current temperature as well as temperature change from the activity levels of each of the seven response types. The same train/test split on trajectories as outlined above (in General approach to model fitting and testing) was used. Models were fit on train data using Ridge regression and the ridge penalty was set using hold-one-out cross validation. Predictions on test data were subsequently reported.

To test if neural activity could control the transition matrix, the GLMs of the Markov model were fit using predicted neural activity during behavior as input. The GLM was a linear function of activity across the seven medullary response types at the time of the bout, 500 ms and 1000 ms into the past. Three timepoints were chosen as model inputs since temperature changes will only occur while the fish is swimming which is a stochastic process. Effects of type activity on swim mode probability were computed on the test data. Specifically, the activity was used to compute a transition matrix from which steady state probabilities were computed. These were binned into quartiles and the average activity of each type across all three timepoints was reported.

#### Statistics and Exclusion criteria

All reported errors are non-parametric bootstrap standard errors. Bootstrapping was performed across fish. Fish with an average swim frequency below 1 Hz were excluded from gradient experiment analysis. Since cold temperatures were used in some constant temperature experiments, the threshold was lowered for all constant temperature experiments to an average of one bout every 1.5 s. See Supplemtary Table 2 for a count of experiments performed, experiments that passed filtering, bout counts before and after edge filtering and number of trajectories used for model fitting.

## Acknowledgements

Research reported in this publication was supported by the National Institute Of Neurological Disorders And Stroke of the National Institutes of Health under Award Number R01NS123887 to MH and by the Office Of The Director, National Institutes Of Health of the National Institutes of Health under Award Number R24OD037693 to MH. The funders had no role in study design, data collection and analysis, decision to publish, or preparation of the manuscript. MH and KAB received salary from the funders. The content is solely the responsibility of the authors and does not necessarily represent the official views of the National Institutes of Health. We thank Jamie D Costabile for designing and building the gradient chamber and writing the control software. We thank Lindsay Anderson, Danica Matovic and Bradley Cutler for valuable comments on the manuscript.

## Author Contributions

Kaarthik A Balakrishnan: Conceptualization; Investigation; Methodology; Formal Analysis; Software; Writing - Original Draft Creation

Martin Haesemeyer: Conceptualization; Methodology; Data Curation; Formal Analysis; Software; Funding Acquisition; Supervision; Writing - Review & Editing

## Supplemental Figures and Tables

**Figure S1: Extended data related to Figure 1.**
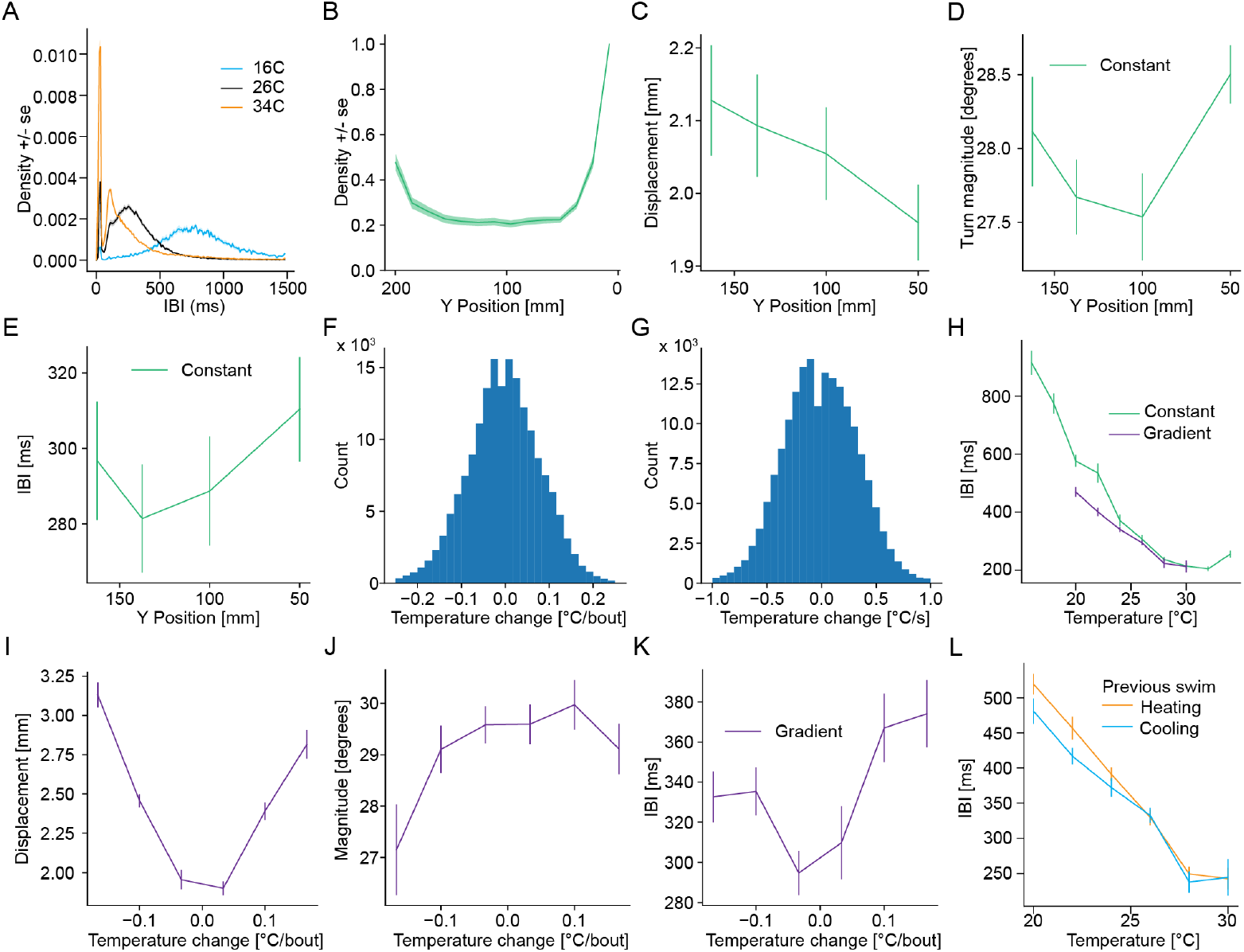
**A** Example distribution of interbout intervals (in ms) at three different temperatures averaged over all the corresponding constant temperature experiments (blue-16 °C, black-26 °C, orange-34 °C). **B** Occupancy of larval zebrafish with respect to y-position averaged across all constant temperature experiments. **C** Median displacement (in mm) of larval zebrafish with respect to y-position, averaged across all constant temperature experiments. **D** Median turn magnitude (in degrees) of larval zebrafish with respect to y-position, averaged across all constant temperature experiments. **E** Median interbout interval (in ms) of larval zebrafish with respect to y-position, averaged across all constant temperature experiments. **F** Histogram of change in temperature experienced by fish per bout in gradient temperature experiments. **G** Histogram of change in temperature experienced by fish per second in gradient temperature experiments. **H** Comparison of median interbout intervals of larval zebrafish at different temperatures for constant temperature (green) and gradient temperature (purple) experiments. **I** Median displacement per bout (in mm) as function of change in temperature in the preceding swim bout, averaged across all temperatures in gradient experiments. **J** Median turn magnitude per bout (in degrees) as function of change in temperature in the preceding swim bout, averaged across all temperatures in gradient experiments. **K** Median interbout interval per bout (in ms) as function of change in temperature in the preceding swim bout, averaged across all temperatures in gradient experiments. **L** Comparison of median interbout intervals (in ms) of larval zebrafish with respect to temperature for different contexts in gradient experiments-orange-heating context, blue-cooling context. All error bars and shaded error regions are bootstrap standard errors across experiments. **Note:** The Y-Position is presented on an inverted axis in all plots to reflect the fact that cold temperatures were always on the side of the chamber with the lowest y-coordinate.

**Figure S2: Extended data related to Figure 3.**
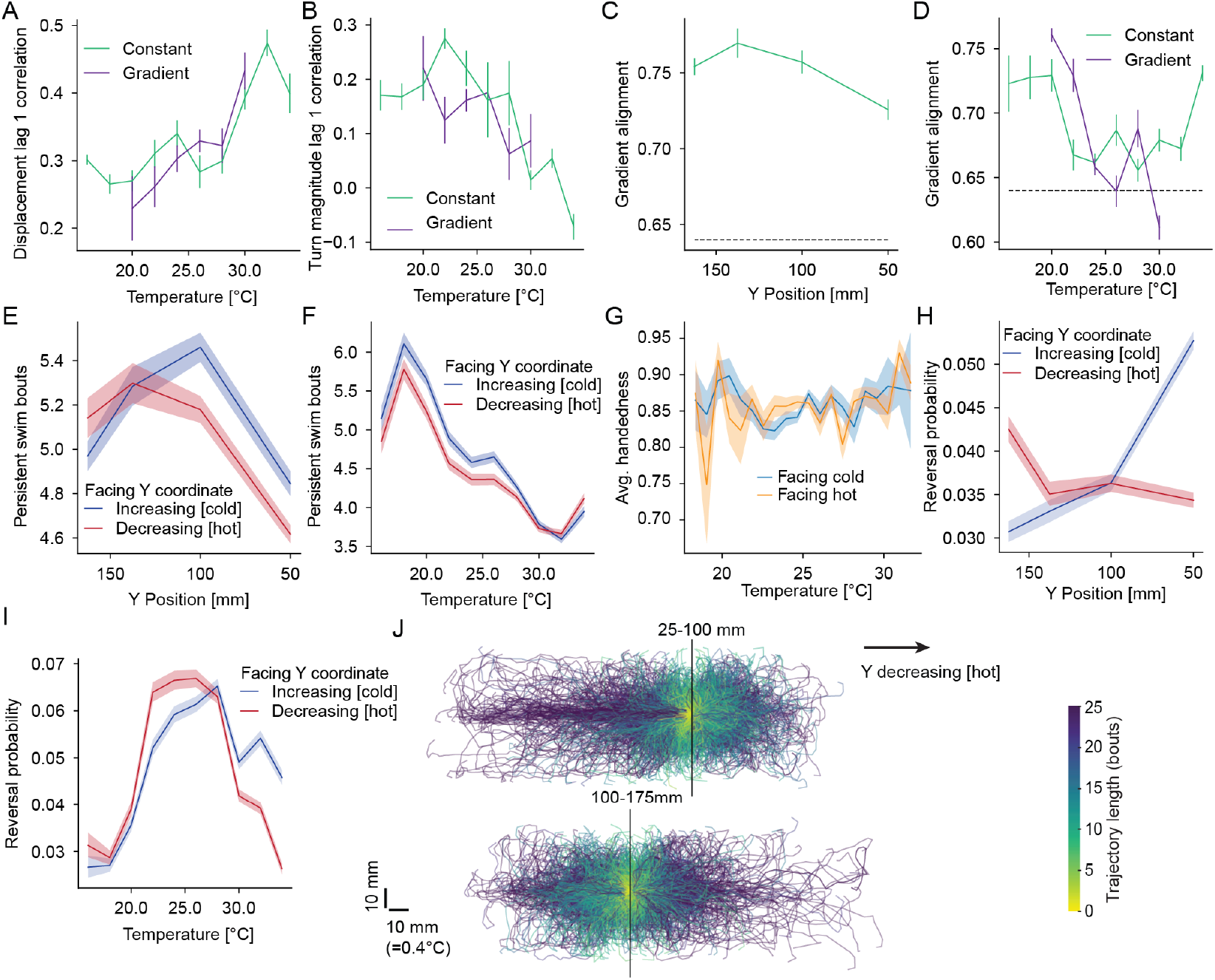
**A** Autocorrelation of displacement in successive bouts with respect to temperature in constant temperature (green) and gradient temperature (purple) experiments. **B** Autocorrelation of turn magnitude in successive bouts with respect to temperature in constant temperature (green) and gradient temperature (purple) experiments. **C** Alignment of larval zebrafish in the direction of the long edge with respect to y-position, averaged across all temperatures in constant temperature experiments. **D** Alignment of larval zebrafish in the direction of the long edge with respect to temperatures, in constant temperature (green) and gradient temperature experiments (purple). **E** Average number of swim bouts in persistent trajectories with respect to y-position under different contexts-facing increasing y coordinates (blue) and facing decreasing y coordinates (red), averaged across all temperatures in constant temperature experiments. **F** Average number of swim bouts in persistent trajectories with respect to temperature under different contexts-facing increasing y coordinates (blue) and facing decreasing y coordinates (red), averaged across all y positions in constant temperature experiments. **G** Fraction of turns in a reversal maneuver that occur in the same direction as the overall reversal with respect to temperature at the start of the maneuver, while facing towards hot temperatures (orange) and cold temperatures (blue). **H** Probability of reversal of trajectories with respect to y position under different contexts-facing increasing y coordinates (blue) and facing decreasing y coordinates (red), averaged across all temperatures in constant temperature experiments. **I** Probability of reversal of trajectories with respect to temperature under different contexts-facing increasing y coordinates (blue) and facing decreasing y coordinates (red), averaged across all y positions in constant temperature experiments. **J** Examples of trajectories until a reversal was initiated from experiments with the larval zebrafish starting at y positions in the chamber (black lines) that correspond to the positions of the temperatures analyzed in Figure 3H; colorbar represents number of swim bouts until reversal start. All error bars and shaded error regions are bootstrap standard errors across experiments. **Note:** The Y-Position is presented on an inverted axis in all plots to reflect the fact that cold temperatures were always on the side of the chamber with the lowest y-coordinate.

**Figure S3: Extended data related to Figure 4.**
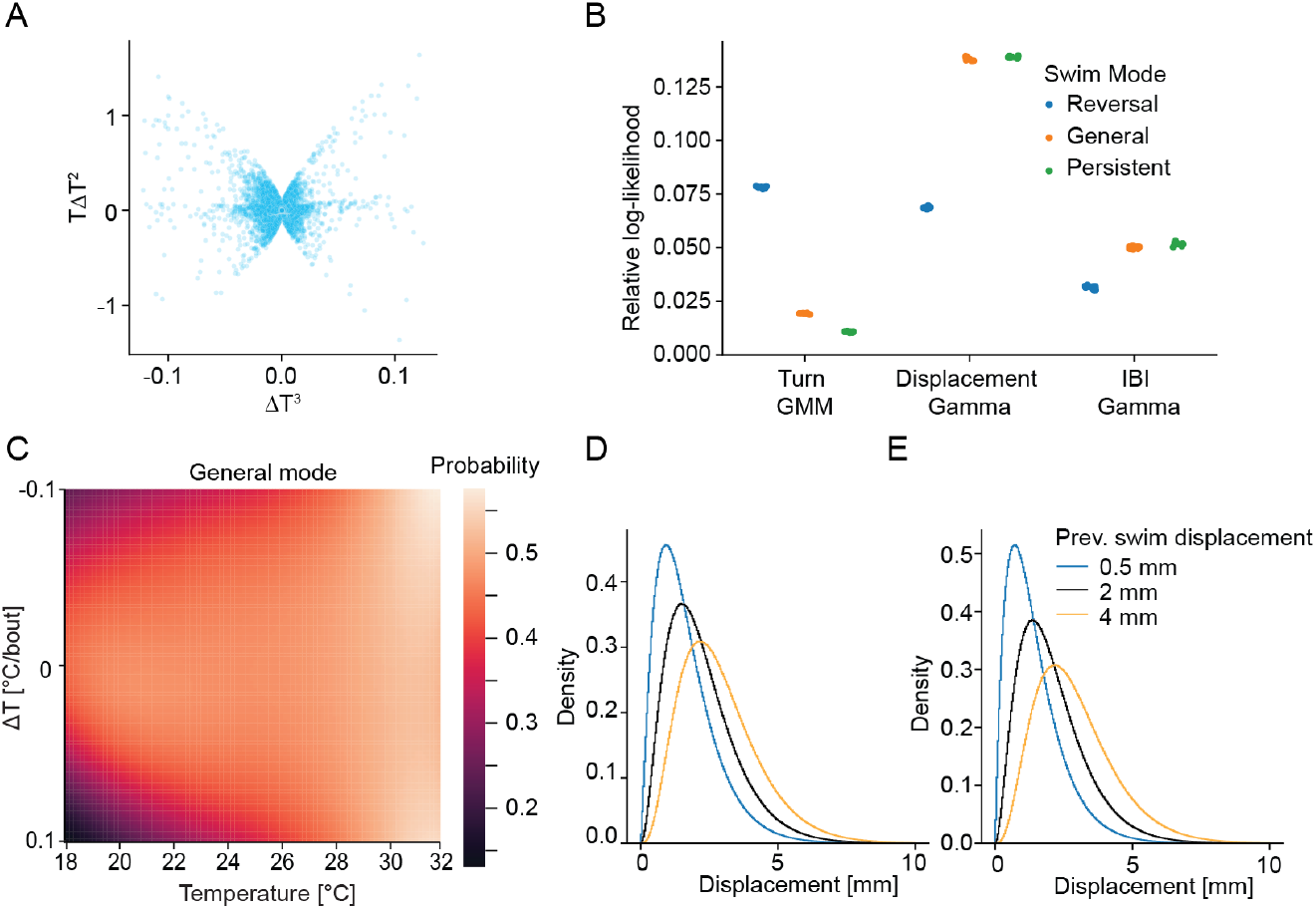
**A** Scatter plot of *T* Δ*T* ^2^ versus Δ*T* ^3^ showing high non-linearity and no correlation across these model inputs (*r* = *−*0.05). **B** Comparison of goodness of fit of emission models for each bout parameter using log-likelihood of model predictions on test-data as an improvement over the chance model which uses the overall data distribution for predictions. Color indicates swim mode. **C** Heatmap of steady state probabilities of the general state depending on temperature and change in temperature for the 3rd order transition model. **D** Dependence of emission of displacement on the displacement of the preceding swim bout in the reversal mode-previous displacement of 0.5 mm (blue), previous displacement of 2 mm (black) and previous displacement of 4 mm (orange). **E** Same as (D) for persistent mode.

**Figure S4: Extended data related to Figure 5.**
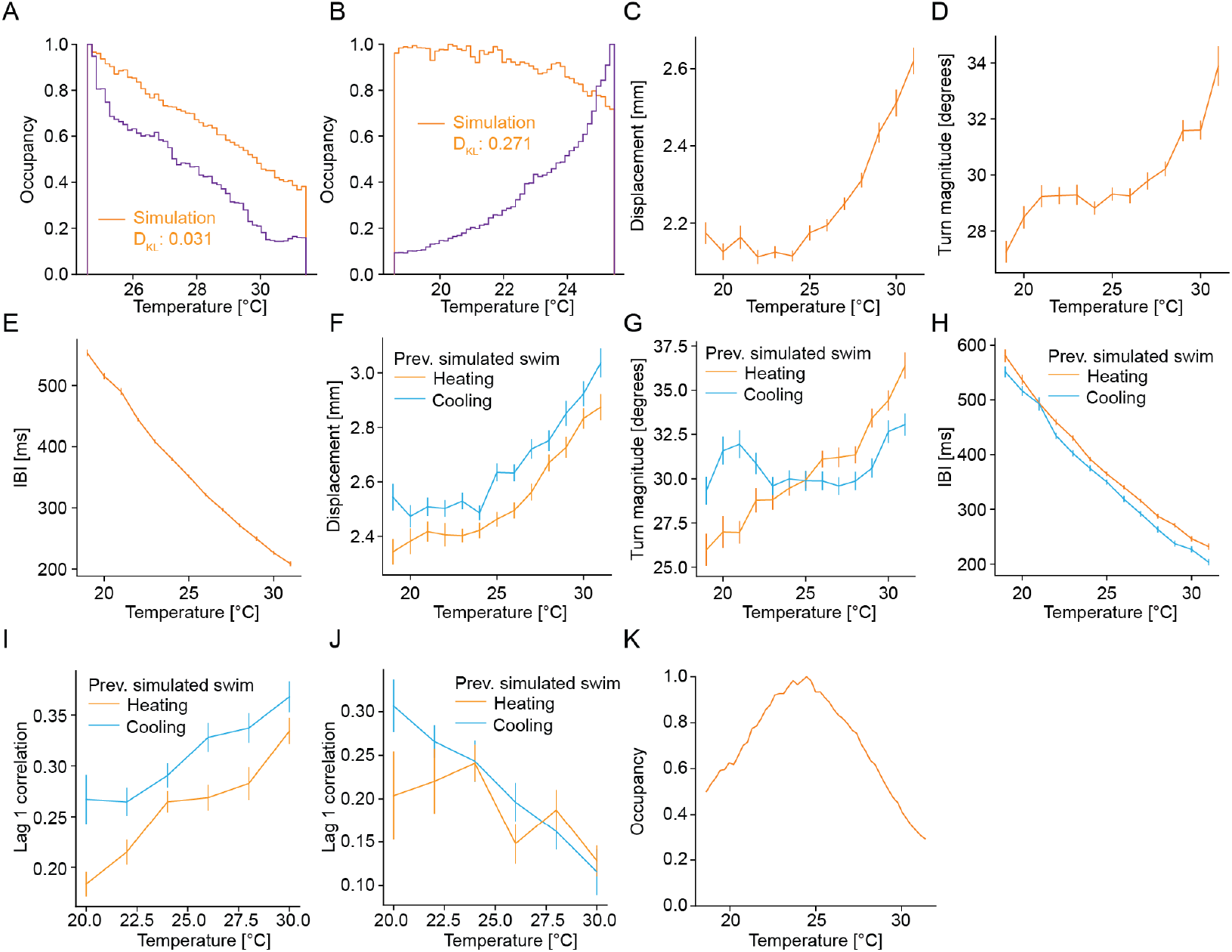
**A** Comparison of occupancy of fish (purple) in hot gradients and simulation using parametric Navigation model for transition model of order 0 (orange). *DKL* inidcates the KL-divergence of the simulation distribution to the experimental distribution. **B** Same as A) but for comparison of occupancy of fish (purple) in cold gradients. **C** Median of generated swim displacement emissions (in mm) for the Navigation model with a 3rd order transition model. **D** Median of generated swim turn magnitude emissions (in degrees) for Navigation model with a 3rd order transition model. **E** Median of generated swim interbout interval emissions (in ms) for the Navigation model with a 3rd order transition model. **F** Comparison of generated median swim displacement emissions with respect to temperature in different contexts-orange-heating context, blue-cooling context for Navigation model with a 3rd order transition model. **G** Comparison of generated median swim turn magnitude emissions with respect to temperature in different contexts-orange-heating context, blue-cooling context for Navigation model with a 3rd order transition model. **H** Comparison of generated median swim interbout interval emissions with respect to temperature in different contexts-orange-heating context, blue-cooling context for Navigation model with a 3rd order transition model. **I** Autocorrelation of successive displacement emissions of 3rd order Navigation model in successive bouts with respect to temperature in different contexts-heating (orange) and cooling (blue). **J** Autocorrelation of successive turn magnitude emissions of 3rd order Navigation model in successive bouts with respect to temperature in different contexts-heating (orange) and cooling (blue). **K** Occupancy of fish simulations using the Navigation model with a 3rd order transition model in a virtual chamber with a temperature gradient from 18 °C to 32 °C. All error bars are bootstrap standard errors across simulations.

**Figure S5: Extended data related to Figure 6.**
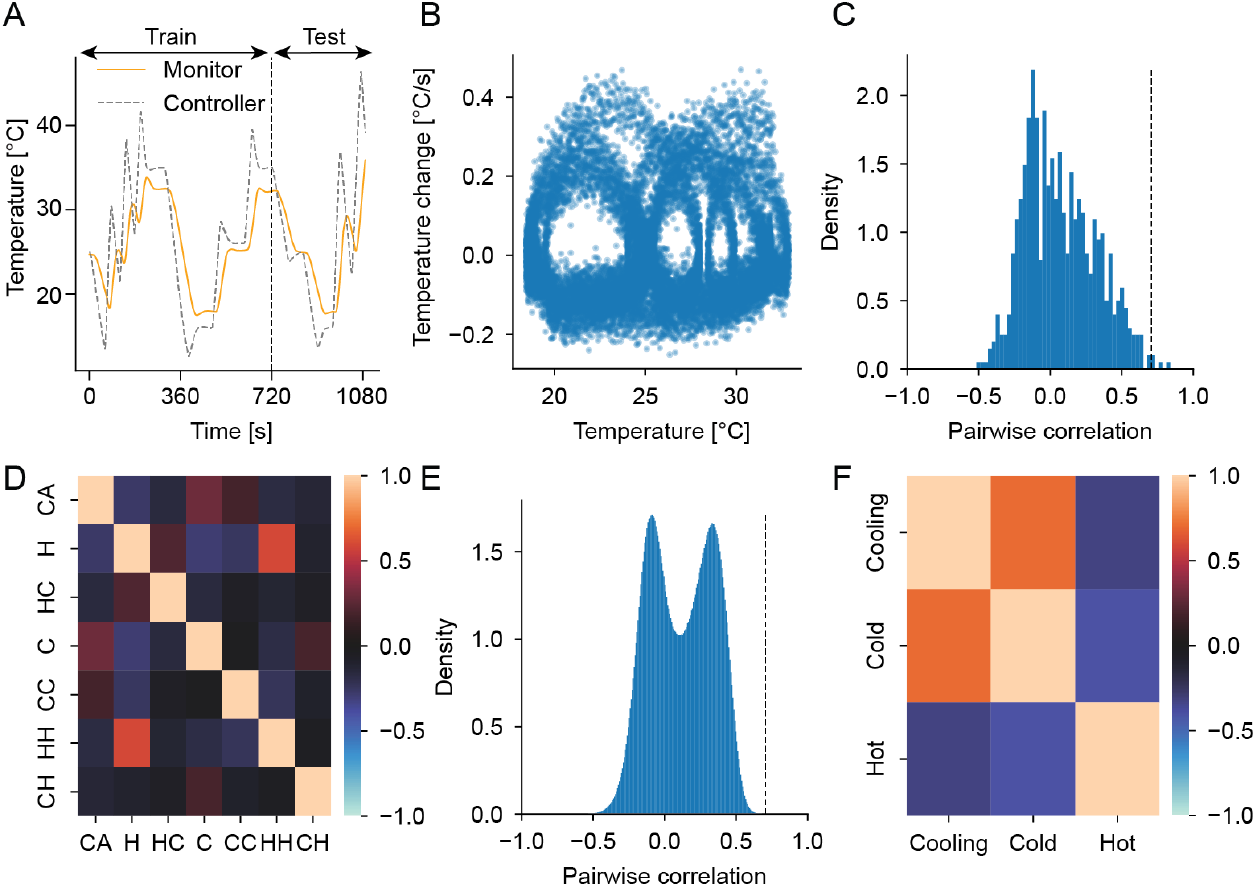
**A** Generated temperature stimulus measured at the controller (black dotted) compared to temperature recorded at the fish using a monitor thermistor (orange). Dashed vertical line indicates split of training versus test data used in MINE fits. **B** Scatter plot of temperature change versus temperature in temperature of the presented stimulus across all experiments. **C** Density of pairwise correlations of unclustered neurons in the medulla. **D** Heat map of pairwise correlations of mean responses between different response types in the medulla-CA: Cold adapting, H: Hot, HC: Hot and Cooling, C: Cold, CC: Cold and Cooling, HH: Hot and Heating, CH: Cold and Heating. **E** Density of pairwise correlations of unclustered neurons in the Trigeminal Ganglion. **F** Heat map of pairwise correlations of mean responses between different response types in the trigeminal ganglion.

**Figure S6: Extended data related to Figure 7.**
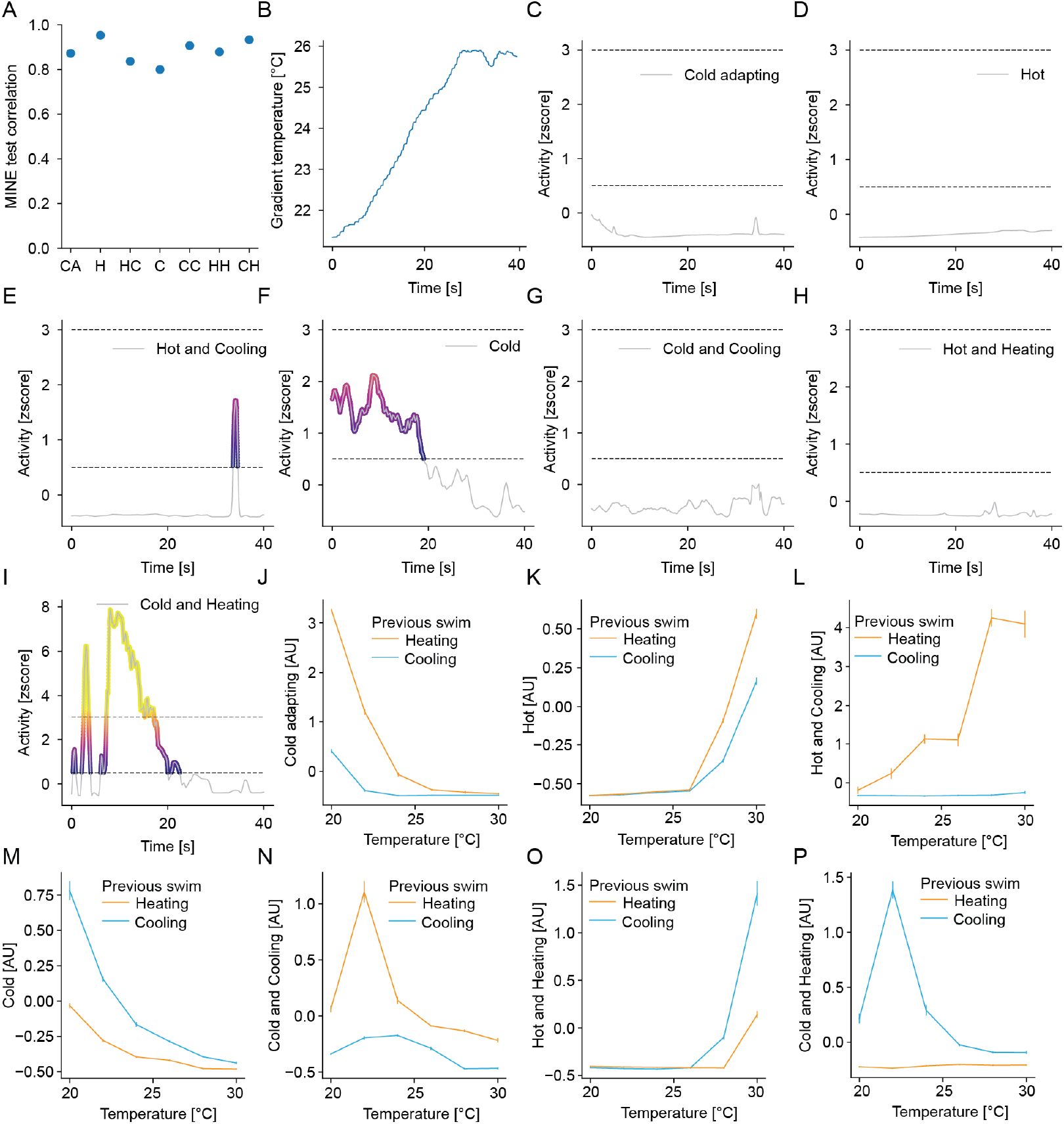
**A** Average test scores of MINE predictions on the deconvolved spike trains for each cluster given 2 seconds of temperature history. **B** Example temperature trace of fish swimming in a gradient. **C-I** MINE predicted neural activity of each cluster for the example trace in (B). The lower dashed line indicates the activity threshold used for coloring the traces in these panels as well as Figure 7A. The upper dashed line indicates the activity level at which the color scale is maxed out at yellow. **J-P** MINE predicted neural activity of each cluster with respect to temperature for different contexts-heating (orange) and cooling (blue).

**Supplementary table 1:** This table contains the averages, 5th percentile and 95th percentile of all draws from the posterior distributions of all model parameters that were fit using STAN.

**Supplementary table 2:** This table contains counts of experiments, fish that passed validations and swimbouts as well as trajectories by experiment type.

